# ProPrep: An Interactive and Instructional Interface for Proper Protein Preparation with AMBER

**DOI:** 10.64898/2026.02.26.708365

**Authors:** Adara Walker, Matthew J. Guberman-Pfeffer

## Abstract

Millions of experimental and AI-predicted protein structures are now available, and the biosynthetic promise of bespoke proteins is increasingly within reach. The functional characterization challenge thus posed cannot be addressed by experimental techniques alone. Molecular dynamics (MD) simulations offer functional screening with atomic resolution, yet accessibility remains limited. Existing computational chemistry software presents stark trade-offs whereby powerful tools require extensive expertise and manual effort, or user-friendly programs function as black boxes that obscure critical preparation decisions. Herein, we present ProPrep, an interactive workflow manager that guides users through expert-quality MD preparation by showing the ‘what, why, and how’ of each step while automating tedious manual operations. Within a single workspace, ProPrep integrates (1) downloading structures from multiple sources (PDB, AlphaFold, AlphaFill), (2) performing homology searches, (3) aligning structures, (4) curating and repairing structural issues, (5) applying mutations, (6) parameterizing specialized residues, (7) converting redox-active sites to forcefield-compatible forms, (8) generating topology and coordinate files, and (9) configuring, executing, and analyzing simulations with active monitoring of key quantities via ASCII visualizations. A key innovation is ProPrep’s extensible transformer framework for detecting, defining, and transforming redox-active sites—including mono- and polynuclear metal centers, organic cofactors, and redox-active amino acids—for forcefield compatibility. We demonstrate the full workflow on a 64-heme cytochrome ‘nanowire’ bundle (PDB: 9YUQ), proceeding from a PDF file to energy minimization of the solvated system (467,635 atoms) for constant pH molecular dynamics—a process demanding 4,819 PDB record modifications and 610 bond definitions—in 18 minutes of user interaction. The entire process is recorded in an interactive session log that can be shared and replayed for reproducibility, making simulation setup a fully transparent process that relies on what was done instead of what was remembered and reported.

## 1. Introduction

Molecular dynamics (MD) simulations offer functional screening at atomic resolution of the millions of experimental and AI-predicted protein structures now available,^1–3^ yet the preparation of a protein structure for simulation remains a surprisingly manual, fragmented, and error-prone process. The raw ingredients—structures, forcefields, solvation models—have matured enormously over the past two decades. Transforming a PDB file into a production-ready topology, however, still demands expertise distributed across multiple software packages, each with its own file formats and input syntax, and where successful completion does not guarantee a physically reasonable result. The problem is not a shortage of tools; it is the gap between their individual capabilities and the integrated, decision-rich workflow that a researcher must execute.

Existing tools impose a trade-off between ease of use and scientific transparency. Automated platforms such as CHARMM-GUI^4^ offer impressive breadth in functionality but, as its developers acknowledge, “contains many pre-determined parameters (e.g., number of steps for minimization or the specifics of how to solvate a protein), which are hidden from users.”^5^ The researcher gains the convenience of downloading scripts at the cost of understanding the choices those scripts encode. Manual preparation with AmberTools^6^ offers more visibility and control, but at the cost of orchestrating file conversions, tracking intermediate outputs, and constructing input scripts by hand. This effort scales poorly and becomes increasingly error-prone with system complexity.

Recent automated pipelines reinforce rather than resolve this trade-off: CHAPERONg^7^ automates GROMACS workflows from structure conversion through trajectory analysis; StreaMD^8^ extends this to high-throughput, distributed GROMACS simulations with cofactor support; drMD^9^ and OpenMMDL^10^ provide analogous automation for OpenMM. All, however, draw the same line around system complexity. drMD, which shares the present work’s emphasis on educational value, transparency, and reproducibility, explicitly excludes proteins with non-natural amino acids, post-translational modifications, or organometallic ligands.^9^ StreaMD integrates MCPB.py^11^ for metal site parameterization but warns that the procedure is “highly system-dependent” and users “are advised to have a thorough understanding” before attempting it,^8^ effectively pushing the preparation burden back onto the researcher.

Most recently, LLM-based agents such as DynaMate^12^ and MDCrow^13^ have pursued fully autonomous MD workflows in which a large language model plans, executes, and self-corrects simulations without user intervention. These represent an intriguing frontier, but their scope is restricted to protein and protein-ligand systems, and their task completion rates vary from 46 to 100% depending on system and model.^12^ Even when a task completes, the resulting simulation may not be physically correct, and such errors are harder to detect precisely because the human has been removed from the decision-making process. Taken together, the existing landscape offers a stark choice: automation that hides decisions, transparency that demands expertise, or AI that bypasses the human altogether. None of these tools spans the full preparation workflow for complex systems while keeping the researcher both informed and in control.

The trade-off is most costly for systems containing redox-active sites. Multi-center redox proteins require oxidation and spin state-specific residue naming, explicit bond directives for metal coordination, and, when redox potentials or electron transfer pathways are of interest, systematic generation of microstate-specific topologies. The practical consequence is that research groups build bespoke automation for their specific enzyme families: CYPWare,^14^ for instance, constructs an end-to-end pipeline for CYP450 heme enzymes using MCPB.py and Marcus theory calculations, but it is not transferable to other metalloprotein classes. For a polymeric multi-heme cytochrome ‘nanowire’ bundle containing 64 bis-histidine-ligated c-type hemes,^15^ building the structure topology from the deposited PDB structure would demand >5,000 PDB record edits and 640 bond definitions *per redox microstate* if every heme was retained. No existing tool addresses this preparation burden at scale for arbitrary redox-active proteins.

Herein, we present ProPrep (Proper Protein Preparation), an interactive workflow manager for AMBER^6^ that automates tedious operations while keeping the researcher informed and in control.

The design philosophy is organized around five principles, collectively ACE IT: *Accessible*, approachable while remaining powerful; *Cohesive*, connected tools sharing a unified workspace with cross-module awareness; *Educational*, explaining the what, why, and how of each preparation step; *Interactive*, involving the user at every decision point; and *Traceable*, transparent decisions recorded in replayable session logs.

The present work describes the program’s architecture (Section 2), capabilities for structure acquisition and comparison (Section 3), structure curation (Section 4), forcefield parameterization of specialized residues (Section 5), redox site handling (Section 6), topology generation (Section 7), and simulation setup, execution, and analysis (Section 8). An application example demonstrating the full workflow on a 64-heme cytochrome ‘nanowire’ bundle is given in Section 9, and the article concludes in Section 10 with a discussion of the scope and future directions for the ProPrep program.

## 2. Program Architecture

### 2.1. Dual Interface Modes

ProPrep provides two interface modes that accommodate users with different expertise levels and workflow preferences: a guided workflow mode that presents tools stage-by-stage through the preparation process, and a full-menu mode that exposes all tools simultaneously.

The guided workflow mode (Figure 1) organizes the preparation process into six sequential stages, displaying only the tools relevant to the current stage. A progress indicator displays stage status, and context-aware suggestions guide users toward recommended next actions based on current workspace contents. Navigation commands allow advancing to the next stage, returning to a previous stage, or skipping stages entirely. This progressive disclosure reduces cognitive load for new users while ensuring logical workflow progression.

**Figure 1.**
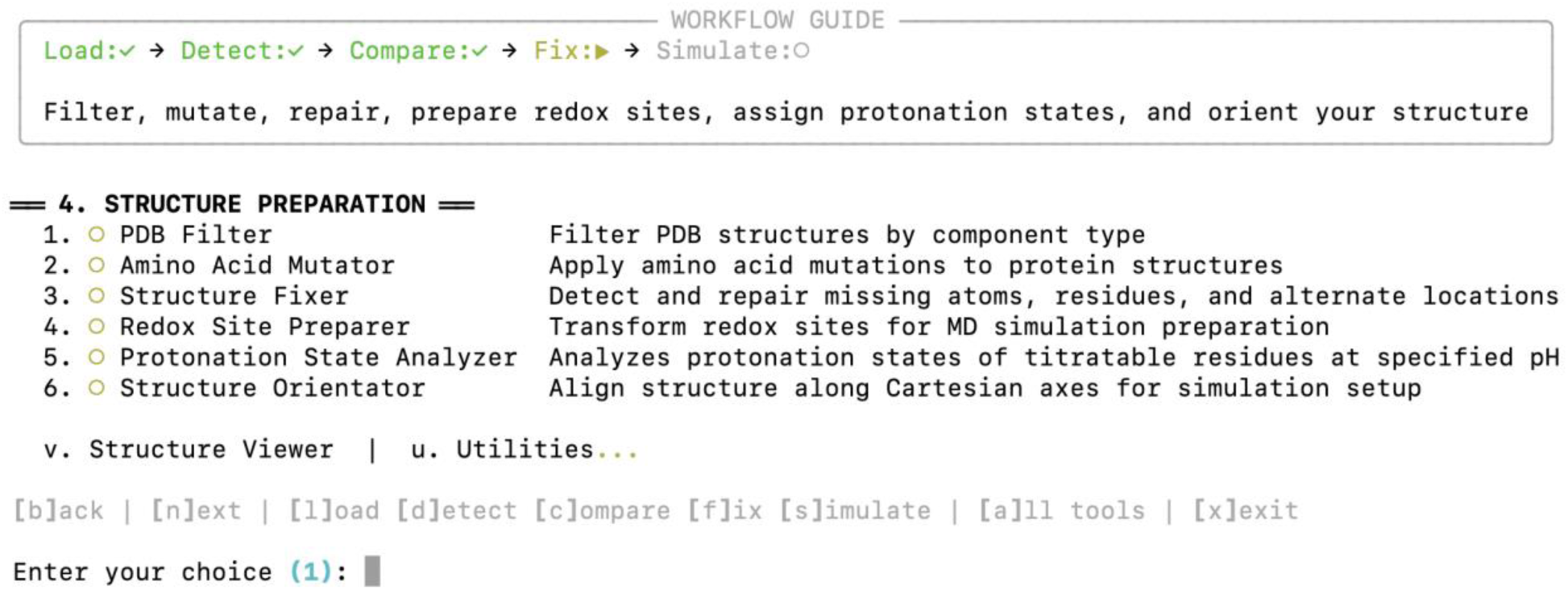
Screenshot of the guided workflow interface in ProPrep. The screen shows the tools available for structure preparation in the Fix stage of the workflow, after the Load, Detect, and Compare stages have either been skipped or completed. The circle/checkmark indicator for each menu option denotes modules that are available given the current state of data in the shared workspace container for the program.

The full-menu mode (Figure 2) exposes all program capabilities simultaneously, organized into numbered sections that mirror the workflow stages. Each tool displays a status indicator showing whether its prerequisites are satisfied (green checkmark) or blocked (yellow circle with explanation of missing requirements), supporting more rapid access by reducing the necessary menu navigations.

**Figure 2.**
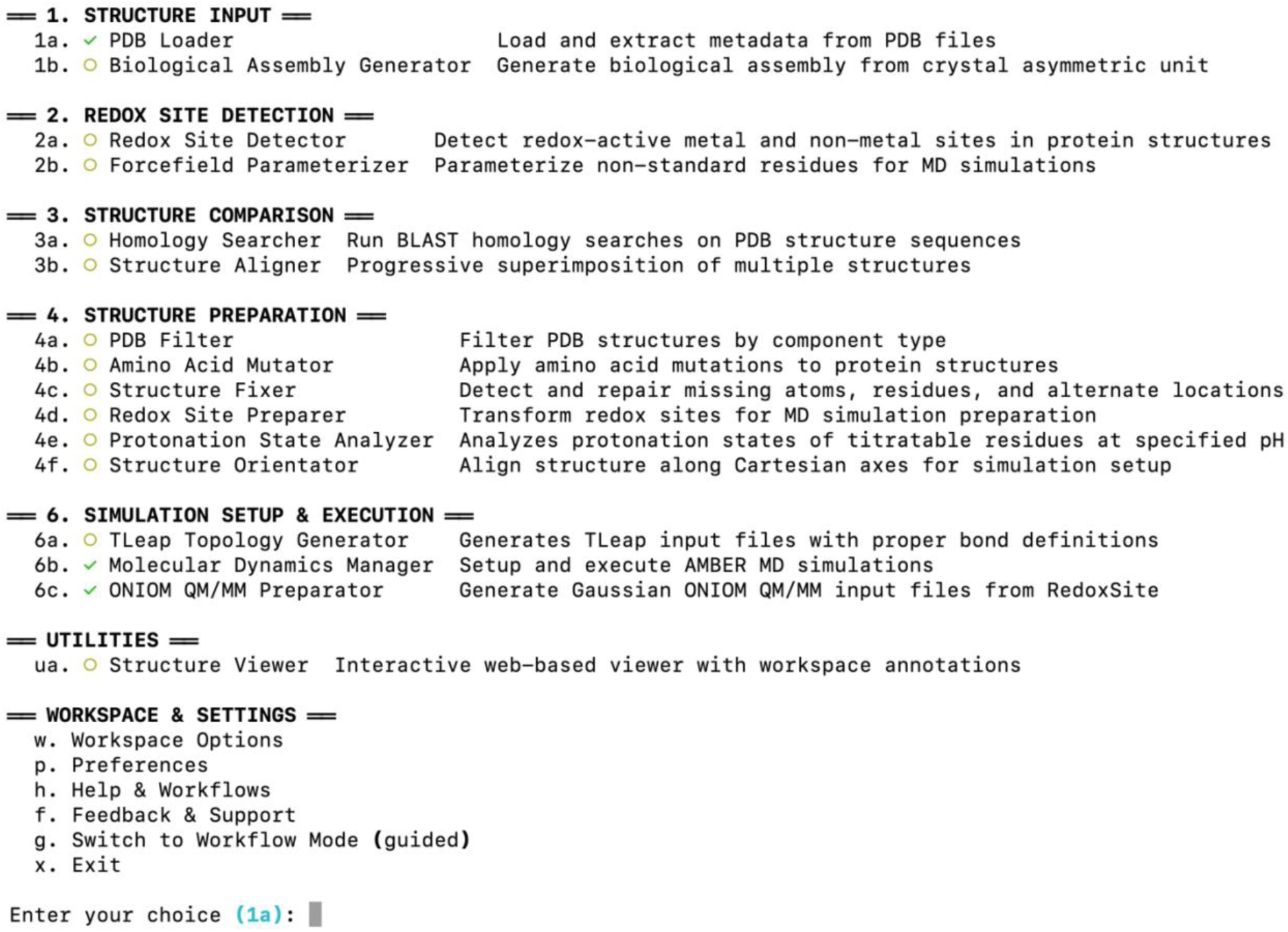
Screenshot of the full menu mode of ProPrep with all the integrated tools exposed and categorized. The circle/checkmark indicator for each menu option denotes modules that are available given the current state of data in the shared workspace container for the program.

Both modes expose the same capabilities and share identical underlying functionality via the shared program workspace. Tool names, stage organization, and navigation patterns are consistent between modes, and users may switch from one mode to the other at any point with full preservation of workspace data.

### 2.2. Workspace System

The workspace functions as a shared data container accessible to all ProPrep modules. When a structure is loaded, parsed metadata, BioPython^16^ structure objects, and file paths are stored under standardized keys. A full inventory of workspace keys in provided in the Supporting information.

The workspace architecture enables inter-module communication that modifies the behavior of downstream functionality in contextually appropriate ways. Five examples illustrate the principle: (1) When the Redox Site Detector defines the atoms and bonds of a redox site, the PDB Filter reads that information to warn users against removing components that participate in redox centers; e.g., a researcher who wishes to retain only chain A is warned against removing chain B if it donates an axial ligand to a heme in chain A. (2) The Protonation State Analyzer reads detected sites to exclude metal-coordinating residues (such as histidines ligating heme iron) from pK_a_ prediction, since their protonation states are constrained by coordination chemistry rather than solvent exposure. (3) Protonation state assignments propagate to the TLEaP Input Generator to ensure that residue naming conventions are forcefield-compliant for the desired protonation state and that the accompanying CPIN file for constant pH molecular dynamics^17,18^ can be successfully generated. (4) The TLEaP Input Generator reads detected sites to construct the bond commands required for AMBER topology generation, translating the site definitions into specific bond directives that enforce correct connectivity. (5) Pending mutations specified in the Amino Acid Mutator are applied by the Structure Fixer—which provides an interface to the MODELLER program^19^ and cleared from the workspace upon success.

The workspace preserves all intermediate results, and modules activate or deactivate menu options based on available data. When a module modifies structure coordinates (e.g., through filtering or repair), it updates the workspace so that downstream modules always access the most current structure.

### 2.3. Interactive Structure Viewer

Visualization is essential throughout the preparation workflow. ProPrep integrates an NGL-based^20^ molecular viewer in the user’s default web browser, available at any point from the main menu. This approach provides interactive 3D graphics without requiring installation of standalone molecular visualization software.

The viewer integrates with ProPrep’s workspace to display contextual annotations that highlight (1) detected redox sites, (2) selected regions when filtering the structure, (3) titratable residues with unusually shifted pKa values, and (4) structures that have been aligned and superimposed. Structures can be shown or hidden independently, and different representation styles (cartoon, surface, ball-and-stick) can be applied to emphasize different features.

### 2.4. Session Recording and Replay

ProPrep records every user interaction with timestamps, creating a log that serves as both documentation and executable history. This session log can be used for three purposes.

#### Resume and Edit

Sessions can be resumed from any recorded interaction. Users browse a table of prior interactions, select a resumption point, and either continue with new interactions or edit the response at that step. Two edit modes are available: smart replay applies subsequent interactions after the modified response, while strict truncate discards subsequent interactions. Either mode enables undoing one or more interactions without losing progress up to that point. The original session is automatically backed up before modification.

#### Replay for Reproducibility

Sessions can be replayed to reproduce exact preparation workflows. Given the same input structure, replay regenerates identical outputs, enabling verification of published results. Session files provide methodological transparency over structure acquisition, curation, preparation, and simulation settings—documentation that is generated in real time and independently of the user’s recollection. Session replay also enables observational learning: trainees can watch recorded sessions from experienced researchers, and instructors can prepare sessions that highlight key decision points.

#### Batch Processing via Templates

Sessions can be converted to reusable templates through a dedicated template creation mode in the session editor. In this mode, the user browses recorded interactions and marks selected responses as variables; for example, marking the PDB ID input as a variable named input_protein. ProPrep auto-detects common variable candidates (structure identifiers, forcefield selections, water models) and pre-populates suggestions that the user can accept, modify, or remove. Each variable is assigned a name, description, and required/optional status. The resulting template file replaces marked responses with placeholders while preserving all other workflow decisions. To apply the template in batch, the session editor generates a CSV file where each row specifies variable values for one run. ProPrep then replays the template once per row, substituting the appropriate values, enabling identical preparation workflows to be applied across homologous proteins or iterative refinements without manual intervention.

### 2.5. Feedback Integration

ProPrep includes integrated issue reporting through GitHub. Users encountering bugs, unexpected behavior, or who have feature requests can submit reports directly from within the program. The submission automatically includes relevant context: Python version, operating system, loaded structure information, and recent session history. This mechanism lowers the barrier to providing feedback and ensures the information needed for diagnosis and resolution is received.

## 3. Structure Acquisition and Comparison

The first stage of simulation preparation involves obtaining and evaluating structural data. ProPrep integrates multiple structure sources—experimental databases, AI-predicted structure repositories, and local files—into a unified loading interface. Complementary modules for homology searching and structure alignment enable comparative analysis without leaving the ProPrep environment or managing intermediate files.

### 3.1. Multi-Source Structure Loading

Protein structures for MD simulations originate from diverse sources with distinct characteristics. Experimental structures from the RCSB Protein Data Bank^1^ provide crystallographic, NMR, or cryo-EM-derived coordinates but may contain missing residues, unresolved side chains, or crystallographic artifacts. AI-predicted structures from AlphaFold^2^ offer complete protein coverage but lack ligands, cofactors, and metal ions essential for many simulations. AlphaFill^21^ addresses this gap by transplanting ligands from homologous experimental structures into AlphaFold predictions. Local files may represent previously prepared structures, homology models, or unpublished experimental data.

ProPrep’s Structure Loader provides a unified interface to all four sources. Structures can be retrieved from the RCSB Protein Data Bank by direct PDB ID entry or through keyword search. The search interface queries RCSB by entry title and returns paginated results displaying PDB ID, title, experimental method, and resolution. Batch downloading of multiple structures is supported. Alternatively, users can search UniProt^22^ by gene or protein name with optional organism filtering.

AlphaFold predicted structures are retrieved by UniProt accession ID or gene name search. ProPrep extracts and displays confidence metrics including mean pLDDT score and quality category. These metrics inform downstream decisions about which regions to trust for simulation and which may require experimental validation.

AlphaFill structures enriched with transplanted ligands and cofactors from homologous structures can be searched and downloaded within ProPrep from the AlphaFill database, enabling simulation of ligand-bound states for proteins lacking experimental holo structures.

Local PDB files can be loaded through an interactive file browser with directory navigation. Files are copied to the project directory to ensure reproducibility and portability. While ProPrep can load mmCIF files for initial inspection, the redox site transformation workflow requires PDB format due to its direct line-editing approach; structures loaded in mmCIF format must be converted before redox site processing.

The Structure Loader also supports loading existing AMBER topology and coordinate files for continuing or analyzing previous simulations with the Molecular Dynamics Manager (Section 8).

### 3.2. Biological Assembly Generation

Crystallographic structures in the PDB often represent an asymmetric unit rather than the biologically relevant oligomeric state. ProPrep provides two routes for obtaining biological assemblies.

When downloading structures from the RCSB PDB with the Structure Loader, users can choose to download a pre-computed biological assembly rather than the asymmetric unit. ProPrep queries the RCSB API for available assemblies and displays them with their oligomeric states. The selected assembly is downloaded directly, with chains relabeled as needed to avoid identifier collisions when symmetry-related copies are merged.

Alternatively, for structures already loaded as asymmetric units, the Biological Assembly Generator module can reconstruct assemblies by applying crystallographic symmetry operations encoded in the PDB header (REMARK 350 records).

### 3.3. Structure Orientation

Membrane protein simulations, surface adsorption studies, and certain enhanced sampling protocols require structures to be oriented along specific Cartesian axes. ProPrep’s Structure Orientor module provides alignment based on geometric analysis of the protein shape. Principal axes or custom vectors can be aligned to user-specified Cartesian directions. Before or after rotation, structures can be centered at the origin using center of mass or center of geometry. The oriented structure is saved with an “_oriented” suffix and automatically becomes the active structure in the workspace.

### 3.4. Metadata Extraction

Experimental PDB files contain extensive metadata concerning structure quality, sequence completeness, biological assembly, disulfide bonds, and heterogeneous groups, *inter alia*. ProPrep parses 29 header record types and 41 REMARK record types, presenting them in categorized menus for targeted inspection. A complete inventory of parsed records is provided in the Supporting Information.

### 3.5. Homology Searching

A target protein may lack an experimental structure in the desired state, or evolutionary aspects of structural features may be of interest. ProPrep integrates NCBI BLAST^23^ searching with direct structure retrieval, removing the need to switch to web-based BLAST services and manually download results. Query sequences can be entered directly, loaded from FASTA files, extracted from chains of loaded PDB structures, or downloaded from RCSB by PDB ID.

Results display as a ranked table showing description, E-value, percent identity, and alignment length. Detailed views provide bit scores, identity counts, and query/subject coordinate ranges. Full pairwise alignments can be examined for selected hits, and results can be exported in XML format or as TSV summary tables.

A key integration feature allows direct downloading of AlphaFold predicted structures for BLAST hits. ProPrep parses hit descriptions for UniProt accessions and retrieves corresponding AlphaFold structures, saving them to a homologs/ subdirectory with confidence metrics. This capability connects sequence homology to structural comparison without manual intervention.

### 3.6. Structure Alignment

Structural superposition enables comparative analysis, quality assessment, and transfer of ligands or cofactors between related structures. ProPrep implements progressive multi-structure alignment with special handling for non-protein components.

In the alignment workflow, the user selects structures from the workspace, a local directory, or directly downloads structures from the RCSB PDB. One structure is designated as the fixed reference; the others are transformed to superimpose onto it. The user defines which residues drive the alignment calculation, choosing from all common residues (excluding water), specific residue ranges, or residues belonging to detected redox sites for active-site comparisons.

A residue mapping strategy establishes correspondence between reference and target residues: (1) sequence-based mapping via pairwise alignment for homologous proteins, (2) structure-based mapping via TM-align^24^ for proteins sharing a fold but not sequence, (3) manual specification assuming identical numbering, or (4) user-defined correspondences. For sequence-based and manual strategies, the user selects which atoms within each residue (e.g., alpha carbon, backbone, all) contribute to the superposition. The superposition is executed progressively with cumulative residue addition for sequence-based and manual strategies, or by direct TM-align transformation for structure-based alignment.

A notable feature of the Structure Aligner concerns the handling of non-protein residues (ligands, cofactors, ions), which cannot be mapped by sequence alignment. These residues are separated from protein residues before alignment and handled through spatial proximity mapping after a preliminary protein superposition. First, protein residues are aligned to establish the transformation matrix. For each non-protein residue in the reference structure, ProPrep identifies target residues with matching names and compares geometric centers in the aligned coordinate frame. Candidates are presented sorted by distance, and the user selects the corresponding residue or adds the reference non-protein residue to the target structure at its transformed position. This approach enables transfer of ligands, cofactors, or ions from one structure to another—a capability that is particularly useful when combining structural information from multiple sources, such as transplanting calmodulin-bound Ca^2+^ ions from a crystal structure (PDB 1CLL) into an NMR-derived model missing them (PDB 2O60).

Reference and target structures are saved with alignment suffixes, preserving the originals. When multiple targets are aligned, the user selects one for downstream processing while all remain available in the workspace.

## 4. Structure Curation

Experimental structures rarely arrive simulation-ready. Missing residues from disordered regions, unresolved side chain atoms, crystallographic waters of uncertain significance, and unspecified protonation states must all be addressed before topology generation. ProPrep provides integrated tools for structure curation with awareness of detected redox sites, so that repair operations do not inadvertently disrupt catalytically essential components.

### 4.1. Component Filtering

The PDB Filter module enables selective retention of structure components—protein chains, nucleic acids, ligands, cofactors, ions, and water molecules—with analysis methods for distinguishing structurally important waters from bulk solvent.

PDB structures are filtered in a hierarchical order. Users first select which model to process if the structure contains multiple models, then select which chains within that model should be retained. Residues in each selected chain are classified as protein, DNA, RNA, water, or heterogeneous (HETATM) group. The canonical 20 amino acids and 4 DNA/RNA nucleobases are recognized, as well as HOH and WAT residue names for water molecules. Any other residue is considered a HETATM group, and the Chemical Component Dictionary^25^ is queried for its name.

The filtering options for each component within a chain are (1) retain the entire component, (2) select specific residues, or (3) discard the entire component. If one or more residues of the component belong to a detected redox site, ProPrep warns against filtering out those residues and provides two additional options: (4) retain only residues belonging to the redox site, or (5) retain redox site residues and ±N flanking residues.

When filtering water molecules, ProPrep provides several analysis methods to assess structural importance. These tools, detailed in the Supporting Information, include assessing (1) distance from metal ions, (2) participation in hydrogen-bonding networks, (3) B-factors, (4) burial within the protein, and (5) proximity to protein-protein interfaces. Users select which methods to apply, optionally adjust parameters, view results as a color-coded table, and select which waters to retain.

### 4.2. *In Silico* Mutagenesis

The Amino Acid Mutator module displays the protein sequence and allows the user to specify which proteogenic residue(s) should be mutated to a standard or modified amino acid. Mutations to a canonical amino acid are applied using MODELLER in the Structure Fixer module: the backbone atoms are preserved, the side chain is replaced, and energy minimization resolves any clashes. For mutations to a modified amino acid, users select which atoms to retain from the original residue (typically backbone atoms N, CA, C, O; optionally CB), and the TLEaP Topology Generator builds missing atoms based on the target residue’s forcefield definition. This approach requires that the modified residue has forcefield parameters available, which can be obtained using the Forcefield Parameterizer module (Section 5). ProPrep’s workspace seamlessly handles the communication between the Amino Acid Mutator, Structure Fixer, Forcefield Parameterizer, and TLEaP Topology Generator.

### 4.3. Structure Repair

The Structure Fixer module detects missing residues, missing atoms, and residues with alternate locations. Missing residues are detected by analyzing REMARK 465 or SEQRES metadata records of a PDB file, comparing the deposited FASTA sequence to the ATOM records, or by detecting discontinuities in residue numbering. Missing atoms are detected by parsing REMARK 470 records or by comparing to the Chemical Component Dictionary template for each unique residue type. Alternate locations for residues with multiple conformations are identified via altloc identifiers or partial occupancy values.

Options to resolve these issues are then provided. Each segment of missing residues can be built—in part or in full—via MODELLER, capped with an acetyl (ACE) or N-methylamide (NME) group, or left unrepaired. Missing atoms are added by the TLEaP Topology Generator at a later step. Alternate locations are presented for each residue with their relative occupancies, and the user interactively selects which conformation to retain.

The structural repair process can change chain identifiers, residue identifiers, and Cartesian coordinates. ProPrep maintains a ResidueMapper that tracks atoms through the process and synchronizes any previously detected redox site definitions with the final structure. This synchronization allows bond directives to be auto-generated with the correct residue numbering for a repaired structure by the TLEaP Topology Generator.

### 4.4. Protonation State Analysis

Correct protonation states are essential for accurate electrostatics in MD simulations. The Protonation State Analyzer predicts the protonation state of selected residues at a target pH and assigns forcefield residue names for those states. For example, an ASP residue can be renamed ASH if it should be protonated or AS4 if it should be titrated in constant pH molecular dynamics. Terminal residues are excluded when assigning titratable residue names because parameters are not available in the standard AMBER forcefields. Users are also given the option to exclude residues that are part of defined redox sites from protonation assessment—a necessary precaution to avoid spurious pKa predictions due to metal coordination.

The module computes protonation probabilities using the Henderson-Hasselbalch equation with PROPKA-predicted^26^ pKa values. Residues are classified as protonated or deprotonated based on a user-selected probability threshold. A results table displays chain IDs, residue IDs, pKa values, shifts (ΔpKa) from textbook reference values, protonation probabilities, and expected charges. The user can launch the Structure Viewer to visualize specific residues and their immediate environments to interpret the predicted pKa shifts.

The analyzer generates pH titration profiles for a user-specified pH range, displayed as ASCII plots directly in the terminal. Users can query by a specific pH or charge, and ProPrep will output the corresponding data. Profiles can be exported to CSV for further analysis or publication-quality plotting.

The analyzer also computes the ideal charge capacitance (∂Q/∂pH) as a function of pH, summed over all titratable residues using the derivative of the Henderson-Hasselbalch equation. The capacitance profile reveals pH ranges where the protein’s net charge changes most rapidly, corresponding to regions where multiple residues have pKa values near the target pH. High capacitance at a given pH indicates that even modest perturbations to the electrostatic environment (e.g., from reduction at a metal active site) would shift the protonation equilibrium of multiple residues simultaneously, with potentially significant functional consequences. This analysis helps select appropriate simulation pH values and interpret charge-dependent behavior.

## 5. Specialized Residue Forcefield Parameterization

Small molecules, modified amino acids, and metal coordination sites generally require parameterization beyond the standard AMBER protein forcefields. The Forcefield Parameterizer module provides wizard-style interfaces to established parameterization protocols for each of these categories. ProPrep integrates residue classification, parameterization, and topology generation into a unified workspace, eliminating the manual file management and format conversions that these multi-step protocols typically demand. Each parameterization workflow is presented through a checklist interface with visual progress tracking; the workflow state is saved to disk after each step, enabling the user to exit and resume in a later session. This capability is essential because the parameterization protocols described below require external quantum chemistry calculations that may run for extended periods on high-performance computing resources.

### 5.1. Residue Classification

Before parameterization begins, the structure must be analyzed to determine which residues require specialized treatment. The Forcefield Parameterizer first collects non-standard residues from two sources: residues associated with detected redox sites (Section 6) are extracted from the workspace with their pre-existing classifications from the detection process, and the remaining structure is scanned for any residue not in the standard amino acid set.

Both sets of specialized residues are then merged and classified through a six-level priority cascade, where the first matching level determines the classification: (1) user-defined classifications take highest priority, enabling manual overrides; (2) redox site classifications are honored directly, routing organometallic cofactors to the Metal Site Parameterizer and organic cofactors to the Small Molecule Parameterizer; (3) single-atom residues whose names match elements in the periodic table are classified as metal ions; (4) a curated mapping of over 200 post-translational modifications and a Chemical Component Dictionary parent-residue lookup identify modified amino acids, which are then routed to the Modified Amino Acid Parameterizer; (5) multi-atom residues classified as NON-POLYMER in the Chemical Component Dictionary and falling within a configurable atom count range are classified as small molecules, with an additional check that reclassifies any small molecule containing a metal atom as a metal site; and (6) residues that pass none of these tests are marked as unknown and require manual classification.

The classification results are displayed in a summary table and stored in the workspace. Users may reclassify any residue before proceeding to parameterization.

### 5.2. Small Molecule Parameterization

Organic small molecules and the organic portions of organometallic cofactors (when parameters for the complete metal-coordinated complex are not yet available) are parameterized using the General AMBER Forcefield (GAFF or GAFF2)^27^ with restrained electrostatic potential (RESP)^28^ partial charges. The workflow guides the user through the standard AMBER parameterization protocol in a step-by-step sequence.

The molecule is extracted from the structure as an isolated residue, and hydrogen atoms are added if absent using the reduce program^29^. ProPrep then generates a two-job Gaussian^30^ input file: the first job performs geometry optimization with frequency analysis for an equilibrium structure, and the second computes the molecular electrostatic potential for RESP charge fitting. Default computational settings follow common practice (B3LYP/6-31+G(d) for optimization; HF/6-31G(d) for ESP), but the user may customize the model chemistry, memory allocation, and processor count for each job. The user specifies the net molecular charge and multiplicity and runs the Gaussian calculation externally.

Upon completion, the workflow resumes by detecting the Gaussian output file. Antechamber^31^ processes the output to assign GAFF2 atom types and fit RESP charges via a two-stage restrained fitting procedure. The parmchk2 utility then generates an frcmod file containing any bond, angle, or dihedral parameters missing from the GAFF2 parameter set, with penalty scores indicating the reliability of each estimated parameter.

A practical complication arises because antechamber renames atoms sequentially (C1, C2, C3, …), whereas PDB files use deposited atom names that follow chemical nomenclature (C4A, C8A, …). If these names are not reconciled, TLEaP will fail when library file atom names do not match PDB atom names. ProPrep addresses this by generating an atom name mapping file that matches atoms between the original PDB and the antechamber-generated MOL2 file by coordinate identity. This mapping is stored and applied automatically during topology generation.

For parameters with high penalty scores—indicating poor transferability from existing GAFF2 parameters, ProPrep offers optional refinement using the Seminario method^32^ or PES-based dihedral fitting. The Seminario method extracts bond stretching and angle bending force constants directly from the Hessian matrix of the quantum mechanical frequency calculation. For torsional parameters, ProPrep generates Gaussian^30^ input files for potential energy surface scans along selected dihedral angles; upon completion, the paramfit program from AmberTools fits dihedral parameters that reproduce the quantum mechanical energy profile. These refinements update the frcmod file, improving parameter accuracy for the specific molecule.

### 5.3. Modified Amino Acid Parameterization

Modified amino acids (e.g., phosphoserine, para-cyanophenylalanine, methylated lysine) require parameters that are compatible with the AMBER amino acid forcefield framework, including proper backbone connectivity and charge constraints. The parameterization protocol differs from small molecule parameterization because the modified residue must integrate seamlessly into the protein backbone when the topology is generated with TLEaP.

The workflow generates initial structures of the modified amino acid in alpha-helical and beta-sheet backbone conformations using TLEaP, with backbone dihedral angles set by cpptraj^33^.

Gaussian input files are created for PES scans of sidechain and functional group dihedrals while the backbone ϕ and ψ angles are held fixed; as with the small molecule workflow, the user may adjust the computational method, basis set, and resource settings. After these calculations complete externally, ProPrep analyzes the PES results, extracts low-energy structures, and generates electrostatic potential calculations for each conformation.

RESP charges are derived using a multi-conformation fitting approach: electrostatic potentials from multiple conformations are combined in the RESP fitting procedure, producing charges that are appropriate across the conformational landscape rather than biased toward a single geometry. The antechamber program processes the lowest-energy structure to generate an AC (AMBER coordinate) file, and the residuegen utility from AmberTools fits charges with appropriate restraints on backbone atoms to maintain compatibility with the parent amino acid forcefield. Finally, parmchk2 generates any missing bonded parameters, and the resulting library and frcmod files are ready for TLEaP.

### 5.4. Metal Site Parameterization

Metal coordination sites require specialized treatment because their electronic structure involves partially filled d-orbitals, variable oxidation states, and coordination geometries that cannot be described by standard organic forcefield parameters. ProPrep implements a parameterization protocol inspired by the Metal Center Parameter Builder (MCPB.py)^11^ from AmberTools, integrated with the redox site definitions described in Section 6.

Before metal-specific parameterization begins, a preprocessing stage prepares the structure and establishes forcefield assignments for all non-metal atoms. The preprocessor performs a component-level triage of the structure, classifying residues as standard protein, organic small molecules, organometallic cofactors, isolated metal ions, or water. Metal atoms are then temporarily removed, TLEaP generates the topology for all remaining components using the selected protein forcefield and any small molecule parameters from Section 5.2, and the metals are reinserted at their original coordinates. Atom types and partial charges for all non-metal atoms are extracted from the resulting topology file (prmtop) using ParmEd.^6^ This approach treats the prmtop as the authoritative source of forcefield data.

Metal site parameterization then proceeds through four steps, with checkpoints between steps that require external Gaussian calculations. In the standard MCPB.py workflow, this preprocessing stage alone is estimated to require <1 hour per metal site due to manual file editing operations. ProPrep replaces these manual edits with interactive prompts that populate the required files programmatically.

*Step 1: Atom Typing and Model Building.* The redox site definition and atom types extracted during preprocessing are read. Since metal atoms were removed before TLEaP, their charges and spin multiplicities are collected from the user. Each atom in the coordination sphere is classified as a metal center, a direct metal ligand, or a second-shell residue. Metal atoms are assigned systematic type names (M1, M2, …) and coordinating ligand atoms corresponding names (Y1, Y2, …) to ensure unique identification in forcefield parameter files. Terminal residues in the coordination sphere are identified by a multi-method voting classifier and capped with ACE or NME groups for proper quantum mechanical treatment. Two quantum mechanical models are then constructed: a small model containing the metal center, its direct ligands, and nearby residues for bond and angle parameter extraction, and a larger model extending further into the protein environment for RESP charge fitting. Gaussian input files are generated for both models.

*Step 2: Bonded Parameters.* The Gaussian geometry optimization and frequency calculation output for the small model is processed. Bond stretching and angle bending force constants are extracted using the Seminario method, which decomposes the quantum mechanical Hessian matrix into atomic pair interactions and projects the resulting eigenvalues onto bond and angle internal coordinates. ProPrep’s implementation handles a subtle numerical issue: when the 3×3 sub-Hessian between two atoms is asymmetric (mathematically permissible even though the full Hessian is symmetric), the eigenvalues can be complex. This occurs predominantly for weak, elongated coordination bonds. ProPrep retains complex arithmetic throughout the projection calculation and takes the real part only after summation, matching the behavior of the reference MCPB.py implementation. The extracted parameters are written to an frcmod file.

*Step 3: RESP Charges.* The Gaussian electrostatic potential calculation for the large model is processed. RESP charge fitting is performed with backbone atom charges restrained to values extracted from the prmtop during preprocessing, ensuring that the metal site charges are compatible with the surrounding protein environment. The fitting follows a two-stage RESP protocol with appropriate restraint weights.

*Step 4: Forcefield Integration.* The bonded parameters from Step 2 and RESP charges from Step 3 are combined into TLEaP-ready library and frcmod files. These files are automatically incorporated into the topology generation workflow described in Section 7.

For systems with multiple metal sites, parameterization is performed independently or each site. An important enhancement to the standard MCPB.py workflow is that ProPrep assigns atom type names through a global registry: Each new site receives type names that do not collide with any previously parameterized site in the same protein. In the standalone MCPB.py workflow, each site is parameterized in isolation, and the user must manually rename atom types to avoid conflicts when combining parameter files for a multi-site system.

## 6. Redox Site Handling

Redox-active sites present unique challenges for MD simulation preparation. Metal sites (e.g., hemes, iron-sulfur clusters, copper sites), organic cofactors (e.g., flavins, quinones), and redox-active amino acids (e.g., tyrosyl radicals, cysteine disulfides) all require specialized treatment for forcefield compatibility. ProPrep provides interactive workflows for the detection, definition, transformation, and generation of forcefield-compliant topologies for one or more redox microstates. Once the user identifies and classifies a site, ProPrep handles the tedious work of renaming and renumbering atoms or residues and specifying bond directives that would otherwise require extensive and error-prone manual editing.

### 6.1. The Challenge

The preparation burden for multi-redox center proteins scales multiplicatively with the number of sites and the redox microstates of interest. Residue IDs, residue names, and/or atom names of non-standard residues in a PDB file may need to be changed in an oxidation and spin-state specific way for forcefield compatibility. The bonds connecting non-standard residues to one another or to standard residues need to be explicitly defined when building the system’s topology—and the failure to do so will not trigger a warning, yielding an unphysical simulation without any indication of an error.

The scale of the problem is best appreciated with a concrete example. For a bundle of polymeric multi-heme cytochrome ‘nanowires’ that contains 64 bis-histidine-ligated c-type hemes,^15^ each heme site requires 75 atom records to be edited and 10 bond directives to be written, giving a total of >4,800 edits and 640 bond definitions. If the study requires sampling different oxidation states, these PDB edits must be performed anew for each redox microstate because the residue names are oxidation state-specific. The bond directives, by contrast, are reusable because the residue IDs do not change. We return to this example in Section 9 where we show that ProPrep enables a user to go from the deposited PDB to a fully solvated, curated, and mutated system for constant pH MD undergoing minimization in a specific redox microstate with alternately oxidized and reduced hemes in 18 minutes of user interactions.

To make the setup practical for any multi-center redox protein, ProPrep addresses the challenge through four integrated capabilities: (1) an inventory scan that identifies potential redox centers, (2) an interactive site definition builder where users group redox centers and specify bonds, (3) a transformer framework that applies forcefield-specific renaming and renumbering, and (4) batch microstate generation that produces PDB files and TLEaP bond directives for each requested oxidation state pattern.

Figure 3 illustrates the complete redox site handling workflow described in the following subsections. Detection (Phases 1–6) identifies and characterizes potential redox centers, producing site definitions with atoms, bonds, and a look-up table for Cartesian coordinate-based tracking, which is needed to follow atoms through changes in residue ID, residue name, and atom name during transformation. Before transformation can proceed, users verify that forcefield parameters exist for each site type; sites requiring custom parameters are routed to the Forcefield Parameterizer (Section 5). Once parameters are available, the transformation is configured by assigning a transformer to each redox site, automatically mapping residues to new IDs to avoid numbering conflicts, and matching site components to the roles that the transformer will operate on. ProPrep then iterates through requested microstates, applying state-specific PDB modifications and resetting the workspace between iterations.

**Figure 3.**
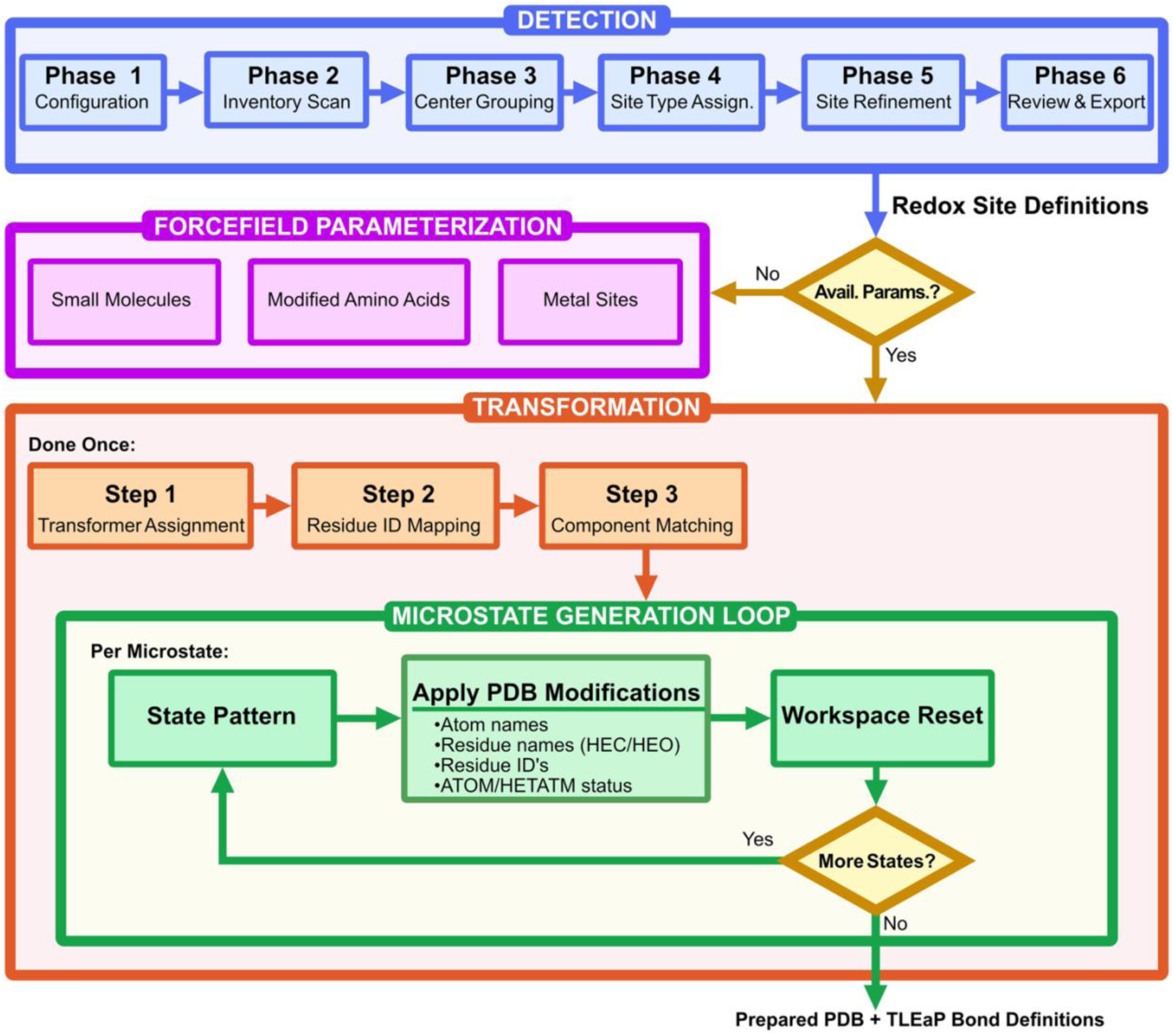
Redox site handling in ProPrep. The flowchart shows the process from detection, to parameterization, PDB ATOM record transformation, and microstate generation. The resulting PDB(s) is/are ready to be processed by TLEaP to build the topology and coordinate files for MD simulations.

### 6.2. Detection

Redox site detection begins with user configuration of search parameters (Phase 1), followed by a structured inventory scan that identifies potential redox-active components in priority order (Phase 2). Any residue classified as an organic cofactor, isolated metal ion, redox-active amino acid, or disulfide-bonded pair is considered a redox center. These centers can then be grouped into redox sites (Phase 3), assigned type classifications (Phase 4), and refined to include all relevant residues with defined bonds (Phase 5). The resulting site definitions are reviewed and exported in Phase 6.

The result of the detection process is a RedoxSite data structure that uses coordinate-based tracking of atoms through changes in atom metadata (residue ID, residue name, atom name) during the later transformation process (Section 6.3). If the structure is modified after detection but before transformation (e.g., to fix missing residues or assign protonation states) ProPrep updates the RedoxSite data object so it remains synchronized with the structure being prepared.

*Phase 1: Configuration.* The configurable parameters are (1) maximum distance for identifying nearby atoms around detected centers (default: 4.0 Å), (2) SG–SG distance for disulfide bond identification (default: 2.5 Å), and (3) which metals and non-metals to include when cataloguing nearby atoms.

*Phase 2: Inventory Scan.* The scan executes four detection passes in priority order: (1) organometallic cofactors—multi-atom non-standard residues containing metal ions (e.g., a heme group is classified as a single organometallic cofactor rather than a metal plus separate ligands); (2) isolated metal ions, identified from a dictionary of 80+ metals; (3) redox-active amino acids (Tyr, Trp, Cys, Met, Phe); and (4) disulfide bonds, detected from SSBOND records or SG–SG distance. Results are displayed in a table listing each detected center with nearby atoms within the search radius, sorted by distance.

*Phase 3: Center Selection and Grouping.* Detected redox centers are selected for further characterization and can be grouped to form multi-center redox sites. Each disulfide-bonded Cys pair is automatically treated as a separate site. If multiple centers share the same residue name, chain, and residue number (e.g., multiple Fe atoms within an Fe4S4 cluster residue), ProPrep offers to group them. Remaining centers require user-specified assignments into sites. This approach handles mono- and polynuclear sites without relying on arbitrary distance criteria.

*Phase 4: Site Type Classification.* Sites can be categorized by type for batch processing. Custom type names (e.g., “bis-his c-type heme,” “Fe4S4 cluster,” “type 1 copper”) serve as labels that enable template-based automation when multiple sites share the same composition.

During Phase 5, the first site of each type is refined manually; ProPrep then applies those refinement settings as a template to subsequent sites of the same type. The settings include (1) how to search for additional residues, (2) which residues to select, and (3) which bonds to define. If a site type name matches an available transformer, that transformer and its options are automatically suggested during preparation (Section 6.3).

*Phase 5: Site Refinement.* Refinement proceeds through an iterative loop: Search → Add residues → Define bonds → Repeat as needed. The starting point for a search can be the geometric center of the current site atoms, the minimum distance from any site atom to each candidate residue, or the minimum distance from a user-specified subset of atoms. Searches use either a distance-based approach (fixed or adaptive radius) or a residue count-based approach (e.g., “find the 2 closest HIS and 2 closest CYS”). Both approaches can be further restricted by element type.

Search results display as a table showing each candidate residue with its name, chain, residue number, atom counts, and distances from the boundary definition. Users select residues to add, with a 3D structure viewer available for visual assessment.

Bond definition follows residue addition. The interface displays residues currently in the site, and users enter residue pairs to bond. Same-residue pairs define intra-residue bonds. For each pair, the interface shows atoms from the first residue and—after selection—displays atoms from the second residue sorted by distance. Multiple target atoms can be selected to define multiple bonds efficiently. If the site type maps to a transformer, the bonds required for forcefield compatibility are displayed for guidance.

*Phase 6: Review and Export.* Completed sites can be reviewed with summary statistics and edited by adding or removing atoms and bonds. Sites can be exported in JSON format (complete data, restorable for future sessions) or as PDB files for visualization.

### 6.3. Site Transformation

The transformer framework converts detected redox sites into forcefield-compatible forms through a five-step workflow. Steps 1–3 execute once to configure the transformation; Steps 4–5 execute per microstate during batch generation (Section 6.4).

*Step 1: Transformer Assignment.* Each site is assigned a transformer—a defined sequence of PDB-editing operations for converting that site type to a forcefield-compatible form. Sites whose type names match available transformers (e.g., heme_bis_his_c_type) receive automatic assignment; others require user selection. Sites already compatible with the target forcefield can use the no_transformation transformer to bypass PDB editing for those residues.

Each transformer specifies (1) the required composition of the redox site, (2) transformation operations for atom metadata, (3) forcefield information (atom types, library files, frcmod files), and (4) available redox/spin states with corresponding residue names. The transformers connect to pre-packaged forcefield parameter sets, and this information is used later in the TLEaP Input Generator to automatically include the appropriate loadamberparams and loadoff commands.

Transformers are pre-packaged for bis-histidine-ligated c-type hemes and disulfide-bonded Cys residues. A pass-through transformer is available for sites that are already forcefield-compatible. A wizard is also provided that guides users in creating custom transformers for their specific redox site of interest. Expanding the library of available transformers is an active direction for development.

*Step 2: Residue ID Mapping.* Transformations that extract atoms into new residues (e.g., heme propionates becoming separate PRN residues) require allocation of new residue IDs. ProPrep analyzes space requirements for all sites collectively, then assigns new IDs to avoid conflicts. This global analysis ensures consistent residue numbering even when multiple sites undergo transformation.

*Step 3: Component Matching.* Each transformer specifies required inputs. For example, the heme_bis_his_c_type transformer requires a heme cofactor, two axial histidines, and two thioether-linked cysteines to be present in the redox site definition. Component matching identifies which residues or atoms in the site fulfill each role. Sites using no_transformation skip this step.

*Step 4: Applying PDB Modifications.* The transformer executes its defined sequence of PDB-editing operations. Each step can consist of the following operations, individually or in combination: (1) atom name changes (e.g., cysteine CB/SG → CBB2/SGB2 when transferred into heme), (2) residue name changes (e.g., HEC → HCR for reduced heme, HCO for oxidized), (3) residue ID assignments (applying the mapping from Step 2), and (4) ATOM/HETATM record status updates.

All modifications are tracked through coordinate-based atom identification, ensuring that bonds and site definitions remain valid even as atom metadata changes.

*Step 5: Resetting the Workspace.* After generating output for a microstate, ProPrep resets the workspace to the pre-transformation state before processing the next microstate. This prevents cumulative metadata changes from corrupting subsequent transformations: each microstate begins from the original detected site definitions.

### 6.4. Microstate Generation

For multi-redox center systems, systematic sampling of oxidation state combinations enables calculation of redox potentials, electron transfer pathways, and cooperativity effects. ProPrep allows any combination of redox/spin states across multiple redox sites to be specified and applies the needed transformations (Steps 4 and 5 from Section 6.3) to produce microstate-specific PDB files that are forcefield-compatible.

Sets of microstates can be specified for batch processing. For example, with the syntax single:1, ProPrep generates microstate PDBs where all sites are in reference state 1, plus separate microstate PDBs where each site is individually varied through all its available states (N+1 microstates for N sites). Full enumeration of states becomes impractical for large N, and ProPrep warns when requested selections exceed 1,000 microstates.

## 7. Topology Generation

With structures curated (Section 4), specialized residues parameterized (Section 5), and redox sites transformed for forcefield compatibility (Section 6), the next stage generates AMBER-compatible topology and coordinate files.

TLEaP, AMBER’s topology-building program, requires carefully constructed input scripts that load forcefields, read structures, define non-standard bonds, add solvent and ions, and save topology/coordinate files. ProPrep generates these scripts through an interactive, guided workflow with full integration of workspace data from preceding modules.

### 7.1. Forcefield and Solvent Configuration

ProPrep presents forcefield options organized by molecular component: protein, nucleic acids, carbohydrates, lipids, small molecules, water, and ions. Each category displays available options with recommendations and compatibility notes. Critical compatibilities are highlighted—for example, that ff19SB^34^ should be used with OPC water rather than TIP3P, since the forcefield was parameterized and validated with OPC^35^.

Users choose between implicit or explicit solvent. For implicit solvent, no solvation section is added to the TLEaP input. For explicit solvent, additional prompts configure box shape, buffer distance, and salt concentration.

ProPrep runs TLEaP with early termination to obtain the water count and system net charge. Using this information, ion counts needed for the target molarity are computed via the SPLIT method.^36^ The methodology is explained in the console output, showing each calculation step with the user’s specific values. Users are advised that TLEaP solvates by tiling a pre-equilibrated water box and removing overlapping molecules, yielding an initial density (∼0.88 g/cc) well below the liquid value. Pressure equilibration compresses the box 10–15% to liquid density, increasing the effective ion concentration by ∼13%.

Users are advised that TLEaP solvates with explicit water molecules at crystallographic

The computed ion counts are placed in two stages for physically reasonable positioning. First, the addions command places neutralizing counterions at positions of lowest electrostatic potential near charged groups. Second, addionsrand places the remaining bulk salt ions at random solvent positions to establish ionic strength.

The TLEaP input template is populated with custom atom type definitions, forcefield parameter loading commands, the structure file path, and bond definitions from the redox site handling workflow (Section 6). The complete TLEaP input file can be viewed and edited before TLEaP is run a second time without interruption to generate the topology.

### 7.2. Multi-Microstate Processing

Processing multiple redox microstates shares the same interactive configuration but adapts for batch efficiency. A shared template with placeholder sections is populated per-microstate with the appropriate custom atom types, forcefield parameters, structure paths, bond definitions, and ion commands based on the ion counts computed for each microstate. This workflow enables processing dozens or hundreds of microstates without manual intervention.

### 7.3. Component Ordering and Validation

Different AmberTools programs expect different component orderings in multi-chain structures. TLEaP requires all protein components (e.g., chains A and B) before the cofactors belonging to those chains, whereas other AmberTools expect all components of chain A before all components of chain B. ProPrep allows the user to reorder the structure appropriately before running TLEaP, and then runs ParmEd to validate and—if needed—reorder atoms to ensure molecules are contiguous.

### 7.4. 12-6-4 Ion Parameters

When the user selects 12-6-4 ion parameters^37^, ProPrep applies ParmEd’s add12_6_4 action to insert LENNARD_JONES_CCOEF (C4 charge-induced dipole) terms into the topology. Rather than using a hardcoded list of ion names, ProPrep scans the topology for single-atom residues to build the AMBER mask dynamically, ensuring all ions in the system are included regardless of naming convention.

The add12_6_4 action requires atomic polarizabilities for every atom type in the topology, but the default lj_1264_pol.dat file covers only standard AMBER types. ProPrep identifies missing atom types, infers their polarizabilities from element identity and hybridization state using values from Miller^38^—the same source as the original file—and writes an augmented polarizability file to the working directory for user inspection and reproducibility. The 12-6-4 potential is automatically skipped for incompatible forcefields (ff19SB, ff15ipq), which renamed atom types in ways that break the polarizability lookup, with a warning explaining that the 12-6-4 parameters were developed and validated with ff14SB^39^.

### 7.5. Constant pH Input Generation

Simulations employing constant pH molecular dynamics require a CPIN file generated by the CPINUTIL.py program from AmberTools. Rather than requiring manual construction of command-line arguments, ProPrep interactively shows which residues in the structure can be titrated, asks the user to select which to include, and allows the user to set initial protonation states. ProPrep then builds the command-line flags and runs CPINUTIL.py on the user’s behalf, but with their desired settings.

## 8. Simulation Setup and Execution

With topology files prepared, the final preparatory stage configures and executes the MD simulation protocol. ProPrep’s Molecular Dynamics Manager handles workflow construction, input file generation, restraint specification, and simulation monitoring through a unified interface. The module is designed around the principle that every simulation parameter should be visible and modifiable, while sensible defaults and educational context support non-specialist users.

### 8.1. Workflow Construction and Template System

MD simulation protocols typically consist of multiple sequential stages—minimization, heating, equilibration with decreasing restraints, and production dynamics—each requiring a separate AMBER input (mdin) file with distinct parameters. ProPrep provides three methods for constructing these multi-stage workflows.

First, built-in workflow presets define complete multi-step protocols validated for common use cases. For example, the standard protein equilibration preset—adapted from the protocol of McCarthy, Ekesan, and York^40^—consists of thirteen steps: initial solvent minimization with the solute restrained, NPT equilibration stages at increasing durations for density convergence, solvent heating from 0 to 300 K, extended NPT equilibration, full-system minimization without restraints, solute heating with positional restraints, and a progressive restraint reduction series (25, 10, 5, and 2 kcal/mol/Å^2^). Each step specifies its input and output coordinate files and its dependencies on prior steps, enabling automatic sequencing.

Second, users may assemble custom workflows from a template library, selecting and ordering individual mdin templates for each stage. Templates can be imported from external files or created through an interactive configuration wizard that guides the user through every namelist parameter group: simulation type, performance options, boundary conditions, restart configuration, simulation length and time step, constraint options (SHAKE/RATTLE), thermostat selection, barostat configuration, nonbonded interaction cutoffs and Ewald summation parameters, output frequency, and advanced options including NMR restraints and special simulation modes. Each parameter is accompanied by a brief explanation of its physical meaning, default value, and guidance on when alternative values are appropriate. This educational metadata is drawn from a curated database of over 100 AMBER parameters organized by category.

Third, users may directly edit mdin file contents through their system text editor for fine-grained control.

Protocols reference templates without copying them; any parameter overrides specified by the user are stored separately and applied when the final mdin files are written.

### 8.2. Restraint Integration

MD restraints are configured through a dedicated Restraint Manager module supporting two categories: positional restraints (restraintmask patterns with force constants) and distance/angle/torsion restraints (AMBER DISANG format with flat-bottom harmonic potentials). Restraints are defined interactively by selecting atoms from the structure, with current geometric values (distances, angles, dihedrals) computed from the coordinates and used as default targets. For systems with redox sites, a restraint mask generator extracts atom identities from the pre-defined redox site definitions and allows the user to select the restraint scope (coordinating atoms only or full ligand residues) and granularity (individual atoms or entire residues), with optional interactive refinement of the selected atoms. Restraints configured in the Restraint Manager are stored in the workspace and automatically detected during simulation setup, where the user assigns them to appropriate workflow steps.

### 8.3. Execution and Monitoring

The configured workflow is organized into a simulation queue with explicit dependency tracking. The user configures each simulation step with an execution engine (pmemd, pmemd.cuda, or sander), MPI task count, and GPU device assignment. For high-performance computing environments, ProPrep generates SLURM job submission scripts with configurable resource allocation (partition, nodes, processors, GPUs, memory, and time limits), module loading commands, and environment setup.

During execution, ProPrep manages workflow dependencies automatically: when a simulation step completes, dependent steps that require its output restart file are started in sequence. The user may monitor running simulations through a real-time interface that parses AMBER output files and displays ASCII-rendered time series of energy, temperature, pressure, volume, and density, along with performance metrics (nanoseconds per day) and estimated time to completion. These terminal-based visualizations require no graphical environment, making them suitable for remote HPC sessions.

Upon completion of each workflow stage, ProPrep performs an automated quality assessment tailored to the simulation type. For minimization, it reports energy convergence, RMS gradient, and whether the energy decreased monotonically. For heating, it verifies that the target temperature was reached and that the temperature ramp proceeded smoothly. For NPT equilibration, it evaluates density convergence, volume stability, and pressure fluctuations relative to system-size-dependent thresholds. For production dynamics, it reports energy conservation and drift statistics. Each assessment includes ASCII plots and summary statistics, providing immediate feedback on whether the simulation proceeded as expected before advancing to the next stage.

### 8.4. Simulation Analysis

ProPrep provides interactive analysis of completed simulations through two complementary approaches: energetics analysis from AMBER output (mdout) files and trajectory analysis from coordinate (nc) files.

#### Energetics Analysis

ProPrep parses mdout files to extract time series of energy components (total, kinetic, potential), temperature, pressure, volume, and density. The simulation type is inferred from the mdin parameters (minimization vs. MD, NVT vs. NPT, heating vs. equilibration), and the user confirms or overrides the detection. Stage-specific quality assessments are then applied: minimization assessments report energy convergence, RMS gradient, and monotonicity of the energy decrease; heating assessments verify that the target temperature was reached and that the ramp proceeded smoothly; NPT equilibration assessments evaluate density convergence by comparing early and late time windows, along with temperature stability, pressure fluctuations, and volume drift; and production assessments report energy conservation and drift statistics. All assessments include ASCII plots and histograms rendered directly in the terminal.

#### Trajectory Analysis

For coordinate trajectory files, ProPrep integrates with pytraj^33^—the Python interface to AMBER’s cpptraj^33^—to provide over 20 analysis methods organized into six categories: (1) structural metrics (RMSD, RMSF, radius of gyration, B-factors), (2) secondary structure and backbone analysis (DSSP, Ramachandran plots), (3) pairwise analyses (contact maps, pairwise RMSD matrices), (4) dynamics (PCA, clustering, autocorrelation), (5) solvation (radial distribution functions, hydration shell populations, solvent-accessible surface area, 3D density maps), and (6) geometric measurements (distances, angles, dihedrals, vector orientations). The user selects analyses from a categorized menu, configures masks and parameters interactively, and receives results as ASCII visualizations with summary statistics. All analysis data can be exported to CSV or JSON for further processing or publication-quality plotting.

## 9. Application Example: Multi-Heme Cytochrome Nanowire Bundle

The preparation burden described in Section 6.1—thousands of PDB edits and hundreds of bond definitions for multi-redox center systems—is not a hypothetical concern. To demonstrate that ProPrep resolves this burden in practice, we prepared a 64-heme cytochrome ‘nanowire’ bundle for constant pH molecular dynamics simulation, proceeding from a raw coordinate file to energy minimization in a single recorded session.

### 9.1. System Description

Our test system is the bundled antiparallel cytochrome ‘nanowire’ from *Desulfuromonas soudanensis* WTL (PDB: 9YUQ),^15^ a halophilic, iron- and electrode-reducing bacterium isolated from deep subsurface brine (Figure 4). The structure was determined by cryo-electron microscopy and comprises 16 protein chains (each 198 residues of the OmcE tetraheme cytochrome), 64 bis-histidine-ligated c-type hemes, and 32 structural Ca²⁺ ions. This system was chosen because it represents an extreme case for MD preparation: the sheer number of redox cofactors, the inter-chain heme ligation (where axial histidine ligands on one chain coordinate to hemes on an adjacent chain), and the need for oxidation state-specific residue naming collectively render manual preparation impractical.

**Figure 4.**
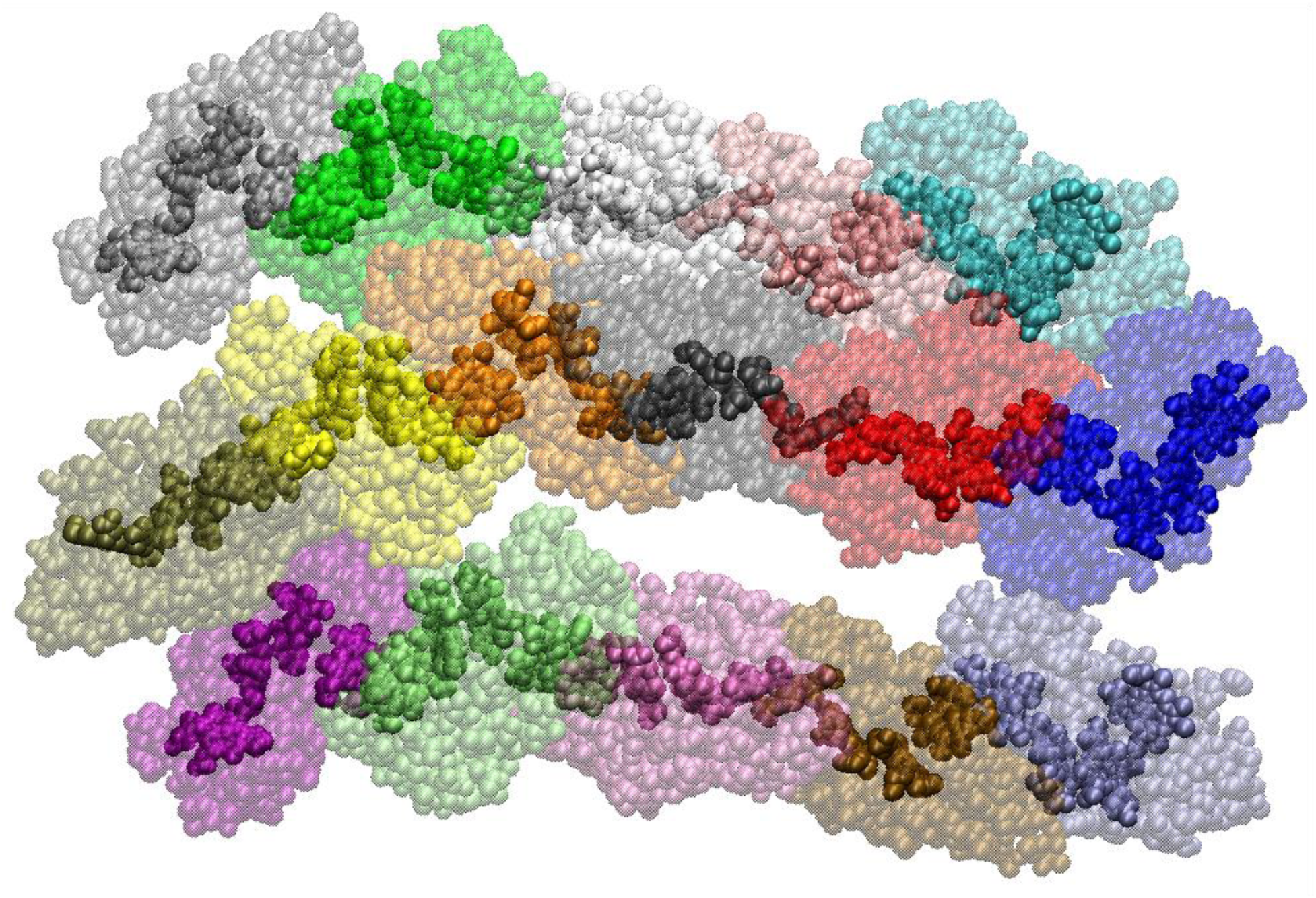
Depiction of the cytochrome ‘nanowire’ bundle from PDB 9YUQ^15^ prepared and minimized for constant pH molecular dynamics using ProPrep after only 18 minutes of user interactions. When solvated with an explicit 12.0 Å water buffer and a 150 mM NaCl salt concentration, the system contains 467,635 atoms.

### 9.2. Workflow Summary

The entire preparation was performed in ProPrep’s guided workflow mode, which organizes the process into sequential stages (Section 2.1). The session comprised 273 user interactions and 66 minutes of wall-clock time from structure loading to simulation launch. Of these 66 minutes, 48 minutes (73%) were consumed by external tool computation—MODELLER for mutagenesis (9 min), ProPrep’s transformer engine processing 61 sites (11 min), PROPKA for pK_a_ prediction (<1 min), and TLEaP for solvation and topology generation (27 min). The remaining 18 minutes represent active user time: the 273 decisions spanning structure loading, redox site detection and configuration, component filtering, mutagenesis, structural repair, redox site transformation, protonation state analysis, forcefield and solvation configuration, restraint definition, and simulation setup.

#### Redox site detection and template automation

ProPrep’s inventory scan identified 96 potential redox centers—64 HEC (c-type heme) organometallic cofactors and 32 Ca²⁺ metal ions. The 64 hemes were selected for further characterization and classified under a single site type (heme_bis_his_c_type), which maps to a pre-packaged transformer (Section 6.3). The first site was refined manually: a count-based search for the two nearest histidine and two nearest cysteine residues located the axial ligands and thioether-bonded cysteines, and six bonds were defined: two Fe–His(N_ɛ_) coordinate bonds for axial ligation, two C–C bonds connecting the propionate groups to the porphyrin macrocycle (necessary because the transformation extracts each propionate into a separate PRN residue that would otherwise be topologically disconnected), and two Cys(C_α_)–Cys(C_β_) bonds (required because the C_β_ and S_γ_ atoms of the thioether-bonded cysteines are transferred into the heme residue definition during transformation). ProPrep then applied this detection template to the remaining 63 sites in a single operation.

In the antiparallel bundle, three filaments each terminate at a subunit whose distal heme would normally receive an axial histidine ligand from the next subunit in the polymer—but the cryo-EM structure captures a finite filament, and no next subunit exists. When the template searched for the two nearest histidines, it located a second histidine approximately 9 Å from the iron center—far beyond coordination distance and clearly not ligated. Because parameters for a penta-coordinate heme are not yet available, and because the ligation state of these terminal hemes is not known experimentally, the three sites were flagged and their redox site definitions deleted. The dynamics of interest lie in the filament interior, not at its termini, making this a physically reasonable simplification. The remaining 61 validated sites each contained 75 atoms and 6 bonds. The manual definition of one site required approximately 2 minutes; propagation to 63 additional sites was practically immediate.

#### Structure curation

All 16 protein chains were retained during component filtering. ProPrep’s redox site-aware filter preserved HETATM groups belonging to defined heme sites while removing the three hemes whose redox site definitions had been deleted in the previous step; these hemes, no longer registered as redox sites, were treated as ordinary HETATM groups and filtered out. A single point mutation (Phe47Tyr in chain N) was specified and queued for application. ProPrep’s structural completeness analysis detected no missing residues, no missing atoms, and no alternate conformations requiring selection; the cryo-EM model was already complete. MODELLER was invoked solely to apply the pending mutation; had any of the aforementioned structural issues been present, they would have been resolved in the same operation.

#### Redox site transformation

Each of the 61 sites was assigned the heme_bis_his_c_type transformer, with alternating reduced (31 sites) and oxidized (30 sites) states to represent a mixed-valence microstate. The transformation modified 79 PDB lines per site across 10 operations: renaming heme residues to oxidation state-specific identifiers (HCR for reduced Fe²⁺, HCO for oxidized Fe³⁺), extracting propionate groups into separate PRN residues, transferring thioether-bonded cysteine atoms into the heme residue, and updating axial histidine names (PHR/PHO and DHR/DHO for the proximal and distal ligands, respectively). Across all 61 sites, this amounts to 4,819 PDB line modifications performed automatically. Workspace validation confirmed that all 4,575 redox site atoms, 61 centers, and 366 bonds remained consistent with the transformed structure, a critical check, because the bond directives that ProPrep writes into the TLEaP input must reference the correct post-transformation residue identifiers and atom names. A single inconsistency here would produce a silent topology creation error or a TLEaP failure.

#### Protonation state analysis and topology generation

All residues belonging to redox sites were excluded from pK_a_ prediction, which is a necessary precaution for metal coordinated residues. PROPKA analysis at pH 7.0 informed titratable residue naming for constant pH MD: 199 aspartate and 32 glutamate residues were assigned titratable names (AS4 and GL4, respectively), all non-coordinating histidines were set to the N_ε_-protonated tautomer (HIE), and a pH titration profile was generated across the full pH range (0–14) and exported for subsequent analysis.

The TLEaP input file was generated with the constant pH forcefield (ff10), TIP3P water, and 12-6-4 ion parameters. ProPrep automatically populated the input with forcefield loading commands for both the reduced and oxidized heme parameter sets, along with 610 bond definitions (122 coordinate, 244 covalent, and 244 peptide backbone bonds) derived from the 61 redox site definitions. The system was solvated in a rectangular box with a 12 Å buffer, and ion counts for 150 mM NaCl were computed via the SPLIT method^36^: 480 Na⁺ and 278 Cl⁻ ions simultaneously neutralized the system charge (Q = −202) and established the target ionic strength. The full calculation, including each step and the user’s specific values, was displayed in the console to make the ion placement methodology transparent. A CPIN file was generated for constant pH simulation with AS4, GL4, and PRN as titratable residue types.

#### Simulation setup

A five-step equilibration protocol was configured with backbone and redox site positional restraints at decreasing force constants (10, 5, 1, and 0.1 kcal/mol Å²) using ProPrep’s restraint mask generator to apply restraints to the protein backbone as well as selected aotms (FE, NA, NB, NC, and ND) of each heme redox site. The first step—energy minimization with sander—was launched and monitored within the ProPrep session.

### 9.3. Assessment

The central result is the contrast between what this preparation demands manually and what ProPrep automates. Without ProPrep, a researcher would need to (1) manually edit approximately 4,819 PDB atom records with oxidation state-specific residue and atom names (or write a custom script to do this and validate it does not make erroneous edits), (2) write 610 bond definitions referencing the correct residue IDs and atom names, (3) construct a TLEaP input file that loads the appropriate forcefield parameters for both oxidation states and includes all bond commands, (4) compute ion counts for charge neutralization at the target ionic strength, and (5) generate a CPIN file with the correct titratable residue selections.

ProPrep reduced the user’s active involvement in these operations to 18 minutes of interactive decision-making. The template-based automation is a decisive feature: defining one heme site’s composition and bonding pattern, then propagating that definition to 63 equivalent sites, compressed what would be hours of repetitive manual editing into seconds. The session log—273 interactions with timestamps, prompts, responses, and module context—provides a complete, replayable record of every decision made during the preparation.

## 10. Conclusion

The present work describes ProPrep, an interactive workflow manager for molecular dynamics simulation preparation with AMBER. The program was designed to resolve the persistent trade-off between powerful but opaque automated tools and transparent but fragmented manual workflows. ProPrep offers a third path: interactive guidance with full visibility into every decision, while sparing the user from the tedious and error-prone operations those decisions require.

Several features of ProPrep merit emphasis. The single-workspace architecture eliminates the need to install, learn, and coordinate multiple applications—a barrier that is non-trivial for researchers and students entering computational biophysics. Comprehensive session recording provides methodological transparency that is generated in real time rather than reconstructed from memory, and session replay enables both reproducibility and observational learning. The redox site handling framework—from detection through transformation to microstate generation—addresses a preparation burden that, for large multi-redox center systems, would otherwise require thousands of manual PDB edits per oxidation state pattern. The 64-heme ‘nanowire’ bundle application (Section 9) illustrates the point concretely: >4,800 PDB modifications and 610 bond definitions were generated from a single template definition in 18 minutes of user interaction.

Several limitations should be noted. The present implementation targets the AMBER ecosystem; users working with other MD engines (GROMACS^41^, NAMD^42^, OpenMM^43^) would need to adapt the topology outputs. The redox site transformer library currently includes bis-histidine-ligated c-type hemes and disulfide-bonded cysteines; while a wizard supports creation of custom transformers, expanding the pre-packaged library to cover common site types (e.g., iron-sulfur clusters, copper sites, flavins) is an active development priority. The PDB format requirement for redox site transformation precludes direct processing of mmCIF files, though conversion utilities are available. Finally, QM/MM input generation for ONIOM-based calculations on MD-derived configurations is under development and will be described in a future release.

Taken together, ProPrep makes expert-quality MD preparation accessible, reproducible, and instructional.

## Supporting information

Supplementary Data: Session Log for Applicaiton Example

## Acknowledgments

We gratefully acknowledge support for this research through an Institutional Development Award (IDeA) from the National Institute of General Medical Sciences of the National Institutes of Health under grant number P20GM103451. We also acknowledge useful discussions with contributions to the software architecture by software engineer John Cairns and useful discussions with Andrew Smith of the Beratan Lab (Duke University), We also wish to thank Prof. Fengbin Wang for early access to the PDB deposition 9YUQ reported in Ref. 15.

## Author Contributions

A.W. provided software design suggestions, alpha-tested the program, and helped draft the manuscript. M. J. G.-P. developed the program and wrote the manuscript.

## Data and Code Availability

ProPrep source code and documentation will be made freely available upon publication. The session log for the application example (Section 9) is provided as Supporting Information.

