## Supplementary Methods for "ProPrep: An Interactive and Instructional Interface for Proper Protein Preparation with AMBER"

**Supporting Information: ProPrep Workspace Key Inventory**

**Table S1.** Complete inventory of workspace keys in ProPrep. Each entry lists the key name, the module(s) that produce it, the module(s) that require it as input, the data type, and a brief description. Keys prefixed with an underscore are internal and not intended for direct user access. A dash (–) in the "Required By" column indicates that the key is not a declared prerequisite for any module, though it may still be consumed programmatically.

| Key | Produced By | Required By | Type | Description |
|---|---|---|---|---|
| _active_tleap_input_file | TLeap Topology Generator | - | str (path) | Currently active tLEaP input file (internal) |
| aligned_ref_pdb_file | Structure Aligner | Protonation State Analyzer, Amino Acid Mutator | str (path) | Path to aligned reference PDB file |
| aligned_ref_structure | Structure Aligner | - | Structure | Aligned reference Structure object |
| aligned_structures | Structure Aligner | - | list | All aligned structures |
| aligned_target_pdb_file | Structure Aligner | Protonation State Analyzer, Amino Acid Mutator | str (path) | Path to aligned target PDB file |
| aligned_target_structure | Structure Aligner | - | Structure | Aligned target Structure object |
| alignment_residues | Structure Aligner | - | list | Residue-level alignment data |
| alignment_results | Structure Aligner | - | dict | Structural alignment results (RMSD, etc.) |
| alphafill_metadata | Structure Loader | - | dict | AlphaFill metadata |
| alphafill_pdb_file | Structure Loader | Homology Searcher, Protonation State Analyzer, Structure Aligner, Structure Viewer, Redox Site | str (path) | Path to AlphaFill structure with transplanted ligands |

| | | | | |
|---|---|---|---|---|
| | | Detector, Amino Acid Mutator, Forcefield Parameterizer | | |
| alphafill_structure | Structure Loader | - | Structure | BioPython Structure object from AlphaFill |
| alphafill_transplants | Structure Loader | - | list | List of transplanted ligands/cofactors |
| alphafill_uniprot_id | Structure Loader | - | str | UniProt ID used for AlphaFill retrieval |
| alphafold_confidence | Structure Loader | - | dict | pLDDT scores and confidence metrics |
| alphafold_homolog_pdb_file | Homology Searcher | PDB Filter, Protonation State Analyzer, Structure Aligner, Amino Acid Mutator, Forcefield Parameterizer | str (path) | Path to selected AlphaFold homolog PDB |
| alphafold_homolog_structure | Homology Searcher | - | Structure | BioPython Structure for AlphaFold homolog |
| alphafold_homologs | Homology Searcher | - | list | AlphaFold homolog entries |
| alphafold_pdb_file | Structure Loader | Homology Searcher, PDB Filter, Protonation State Analyzer, Structure Aligner, Structure Viewer, Redox Site Detector, Amino Acid Mutator, | str (path) | Path to AlphaFold Database structure |

| | | | | |
|---|---|---|---|---|
| | | Forcefield Parameterizer | | |
| alphafold_structure | Structure Loader | - | Structure | BioPython Structure object from AlphaFold |
| alphafold_uniprot_id | Structure Loader | - | str | UniProt ID used for AlphaFold retrieval |
| altloc_processing_results | AltLoc Selector | - | dict | AltLoc processing results (legacy) |
| biological_assembly_metadata | Structure Loader | - | dict | Biological assembly metadata |
| biological_assembly_pdb_file | Biological Assembly Generator, Structure Loader | - | str (path) | Path to biological assembly PDB file |
| biological_assembly_structure | Structure Loader | - | Structure | BioPython Structure for biological assembly |
| blast_raw_results | Homology Searcher | - | dict | Raw BLAST XML results |
| blast_results | Homology Searcher | - | list | BLAST search results |
| capacitance_per_group | Protonation State Analyzer | - | dict | Per-group capacitance values |
| capacitance_profile | Protonation State Analyzer | - | dict | Capacitance profile data |
| combined_tleap_commands | TLeap Topology Generator | - | list[str] | Combined tLEaP commands for topology generation |
| completeness_results | Structure Fixer | - | dict | Structure completeness analysis results |
| constant_ph_data | Protonation State Analyzer | - | dict | Constant-pH simulation configuration data |

| | | | | |
|---|---|---|---|---|
| constant_ph_residues | Protonation State Analyzer | - | list | Residues selected for constant-pH MD |
| coordinates | - | Molecular Dynamics Manager | str (path) | Path to coordinate file (required by MD Manager) |
| cpin_config | TLeap Topology Generator | - | dict | Constant-pH input configuration |
| cpin_file | TLeap Topology Generator | - | str (path) | Path to cpinutil-generated CPin file |
| detected_redox_sites | TLeap Topology Generator, PDB Filter, Redox Site Detector, Structure Fixer, Redox Site Preparer, Structure Preprocessor (pipeline) | Redox Site Preparer | list[dict] | Detected redox-active sites with metadata |
| disang_export_results | MD Restraint Manager | - | dict | DISANG export metadata (path, counts, status) |
| disang_file | MD Restraint Manager | - | str (path) | Path to exported DISANG restraint file |
| disulfide_bonds | Disulfide Bond Detector | - | list | Detected disulfide bonds (legacy) |
| disulfide_structure | Disulfide Bond Detector | - | Structure | Structure with disulfide bonds (legacy) |
| disulfide_tleap_commands | Disulfide Bond Detector | - | list[str] | TLEaP bond commands for disulfides (legacy) |
| emboss_alignments | EMBOSS Analysis | - | dict | EMBOSS pairwise alignment results |
| emboss_batch_analysis | EMBOSS Analysis | - | dict | EMBOSS batch analysis results |

| | | | | |
|---|---|---|---|---|
| emboss_motifs | EMBOSS Analysis | - | dict | EMBOSS motif/pattern search results |
| emboss_sequence_analysis | EMBOSS Analysis | - | dict | EMBOSS sequence analysis results (pepstats, etc.) |
| excluded_residues | Redox Site Preparer | - | list | Residues excluded during redox preparation |
| filter_selections | PDB Filter | - | dict | User’s filter selections (chains, models, residues) |
| filtered_pdb_file | PDB Filter | - | str (path) | Path to filtered PDB file |
| filtered_structure | PDB Filter, Structure Fixer | - | Structure | Structure after chain/model/residue filtering |
| generated_microstate_pdbs | Redox Site Preparer | - | list[str] | Paths to generated microstate PDB files |
| generated_microstate_tleap_files | TLeap Topology Generator | - | list[str] | Paths to generated microstate tLEaP files |
| global_atom_registry_data | Forcefield Parameterizer | - | dict | Global atom type registry for consistency |
| gpu_ids | Molecular Dynamics Manager | - | str | GPU device IDs for pmemd.cuda |
| ideal_capacitance | Protonation State Analyzer | - | float | Ideal capacitance from protonation analysis |
| is_hplusplus_structure | PDB Filter | - | bool | Whether structure was protonated by H++ |
| ligand_pdb_file | Structure Preprocessor (pipeline) | - | str (path) | Extracted ligand PDB file for parameterization |
| ligand_resname | Structure Preprocessor (pipeline) | - | str | Residue name of extracted ligand |

| | | | | |
|---|---|---|---|---|
| local_metadata | Structure Loader | - | dict | Metadata extracted from local PDB file |
| local_pdb_file | Structure Loader | Homology Searcher, PDB Filter, Protonation State Analyzer, Structure Aligner, Structure Viewer, Redox Site Detector, Amino Acid Mutator, Structure Fixer, Forcefield Parameterizer | str (path) | Path to locally loaded PDB file |
| local_structure | Structure Loader | - | Structure | BioPython Structure object from local file |
| md_residue_names | Protonation State Analyzer | - | dict | MD-compatible residue name mappings (HIE/HID/HIP, etc.) |
| md_simulation_queue | Molecular Dynamics Manager | - | list | Queue of MD simulation configurations |
| md_structure_pairs | Molecular Dynamics Manager | - | dict | Structure pairs (prmtop/rst7) for MD |
| md_template_assignments | Molecular Dynamics Manager | - | dict | Template-to-structure assignments for MD |
| md_workflows | Molecular Dynamics Manager | - | list | Defined MD workflow sequences |

| | | | | |
|---|---|---|---|---|
| metadata | PDB Loader | - | dict | Current PDB metadata (deprecated, backward compat) |
| metal_sites | Redox Site Preparer | - | list[dict] | Metal site definitions from redox preparation |
| microstate_protonation_results | Protonation State Analyzer | - | dict | Microstate-specific protonation results |
| microstate_tleap_template | TLeap Topology Generator | - | dict | tLEaP input template for microstates |
| mpi_tasks | Molecular Dynamics Manager | - | int | Number of MPI tasks for pmemd.MPI |
| mutations_applied | Structure Fixer | - | list | Standard mutations applied during repair |
| net_charge | Protonation State Analyzer | - | float | Net charge of the system |
| non_standard_residues | Forcefield Parameterizer | - | list | Detected non-standard residues |
| nonstandard_mutations_applied | Structure Fixer | - | list | Non-standard mutations applied during repair |
| oniom_input_file | ONIOM QM/MM Preparator | - | str (path) | Path to Gaussian ONIOM input file |
| oniom_setup | ONIOM QM/MM Preparator | - | dict | ONIOM QM/MM layer and calculation configuration |
| orientation_record | Structure Orientator | - | dict | Transformation record (rotation matrix, parameters) |
| oriented_pdb_file | Structure Orientator | - | str (path) | Path to oriented PDB file |
| original_metadata | PDB Loader | - | dict | Original PDB metadata |

| | | | | |
|---|---|---|---|---|
| original_pdb_file | PDB Loader | - | str (path) | Original PDB file path (canonical name) |
| original_residue_numbering_info | PDB Loader | - | dict | Original residue numbering information |
| original_structure | PDB Loader | - | Structure | Original BioPython Structure object |
| parameterized_residues | Forcefield Parameterizer | - | dict | Parameterized non-standard residue data |
| parm7_file | TLeap Topology Generator | - | str (path) | Path to AMBER topology (.parm7) file |
| pdb2pqr_output_file | Protonation State Analyzer | - | str (path) | Path to PDB2PQR output file |
| pdb2pqr_pqr_file | Protonation State Analyzer | - | str (path) | Path to PQR file from PDB2PQR |
| pdb2pqr_target_ph | Protonation State Analyzer | - | float | Target pH used for PDB2PQR protonation |
| pdb_file | PDB Loader, Structure Fixer | Small Molecule Parameterizer | str (path) | Current PDB file path (deprecated, backward compat) |
| pending_mutations | Amino Acid Mutator, Structure Fixer | - | list | Queued standard amino acid mutations |
| pending_nonstandard_mutations | Amino Acid Mutator, Structure Fixer | - | list | Queued non-standard amino acid mutations |
| pending_parameterizations | Forcefield Parameterizer, Structure Preprocessor (pipeline) | - | list | Queued parameterization tasks |

| | | | | |
|---|---|---|---|---|
| preferred_amber_engine | Molecular Dynamics Manager | - | str | Selected AMBER engine (pmemd.MPI or pmemd.cuda) |
| prepared_pdb | Structure Preprocessor (pipeline) | - | str (path) | Final preprocessed PDB file |
| preprocessing_atom_data | Structure Preprocessor (pipeline) | - | dict | Atom data from preprocessing steps |
| preprocessing_frcmod_files | Structure Preprocessor (pipeline) | - | list[str] | Frcmod files from parameterization |
| preprocessing_isolated_metals | Structure Preprocessor (pipeline) | - | list | Isolated metal ions found in structure |
| preprocessing_lib_files | Structure Preprocessor (pipeline) | - | list[str] | Library files from parameterization |
| preprocessing_metal_free_pdb | Structure Preprocessor (pipeline) | - | str (path) | PDB file with metals removed |
| preprocessing_metal_reinsertion_map | Structure Preprocessor (pipeline) | - | dict | Map for reinserting metals after preprocessing |
| preprocessing_metal_types | Structure Preprocessor (pipeline) | - | dict | Metal type classifications for parameterization |
| preprocessing_modified_aa | Structure Preprocessor (pipeline) | - | list | Selected modified amino acid leaprcs |

| | | | | |
|---|---|---|---|---|
| preprocessing_nonstandard_ff | Structure Preprocessor (pipeline) | - | dict | Non-standard residue force field selections |
| preprocessing_organic_ff | Structure Preprocessor (pipeline) | - | str | Selected organic molecule force field (leaprc) |
| preprocessing_organometallic_ff | Structure Preprocessor (pipeline) | - | str | Selected organometallic force field |
| preprocessing_output_dir | Structure Preprocessor (pipeline) | - | str (path) | Output directory for preprocessing |
| preprocessing_protein_ff | Structure Preprocessor (pipeline) | - | str | Selected protein force field (leaprc) |
| preprocessing_protein_input | Structure Preprocessor (pipeline) | - | str (path) | Protein PDB input for tLEaP |
| preprocessing_residue_sequence_map | Structure Preprocessor (pipeline) | - | dict | Residue-to-sequence mapping after preprocessing |
| preprocessing_triage | Structure Preprocessor (pipeline) | - | dict | Residue triage results (standard/nonstandard/metal) |
| preprocessing_water_model | Structure Preprocessor (pipeline) | - | str | Selected water model (leaprc) |
| processed_structure | AltLoc Selector | - | Structure | Structure after AltLoc selection (legacy) |
| propka_pka_values | Protonation State Analyzer | - | dict | PROPKA-calculated pKa values |

| | | | | |
|---|---|---|---|---|
| protonation_method | Protonation State Analyzer | - | str | Method used for protonation (propka, pdb2pqr, etc.) |
| protonation_pdb_file | Protonation State Analyzer | - | str (path) | Path to protonated PDB file |
| protonation_results | Protonation State Analyzer | - | dict | Protonation state analysis results |
| rcsb_download_info | Structure Loader | - | list[dict] | Download info dicts (pdb_id, path, timestamp) |
| rcsb_metadata | Structure Loader | - | dict | PDB metadata from RCSB |
| rcsb_metadata_list | Structure Loader | - | list[dict] | Metadata for all downloaded structures (batch) |
| rcsb_pdb_file | Structure Loader | Homology Searcher, PDB Filter, Protonation State Analyzer, Structure Aligner, Structure Viewer, Redox Site Detector, Amino Acid Mutator, Structure Fixer, Forcefield Parameterizer | str (path) | Path to PDB file downloaded from RCSB |
| rcsb_pdb_files | Structure Loader | - | list[str] | Paths to all downloaded RCSB PDB files (batch) |
| rcsb_structure | Structure Loader | - | Structure | BioPython Structure object from RCSB |
| rcsb_structures | Structure Loader | - | list[Structure] | All downloaded Structure objects (batch) |
| redox_restraint_info | MD Restraint Manager | - | dict | Redox restraint metadata |

| | | | | |
|---|---|---|---|---|
| redox_restraint_mask | MD Restraint Manager | - | str | AMBER mask for redox-site restraints |
| redox_sites | Structure Preprocessor (pipeline) | - | list[dict] | Synced redox sites after preprocessing |
| redox_transformer_mappings | PDB Filter, Redox Site Detector | - | dict | Mappings for redox state transformers |
| remove_hydrogens_for_md | PDB Filter, Redox Site Detector | - | bool | Flag to remove hydrogens before MD preprocessing |
| repaired_pdb_file | Structure Fixer | - | str (path) | Path to repaired PDB file |
| repaired_structure | Structure Fixer | - | Structure | Structure after missing residue/loop repair |
| residue_numbering_info | PDB Loader | - | dict | Residue numbering info (deprecated, backward compat) |
| restraint_integration_config | Molecular Dynamics Manager | - | dict | Restraint integration configuration for MD |
| restraint_mask_generated | MD Restraint Manager | - | bool | Whether restraint mask was generated |
| restraint_structure_source | MD Restraint Manager | - | str (path) | Source structure for restraint generation |
| restraints | MD Restraint Manager | - | list[dict] | List of restraint definitions |
| rst7_file | TLeap Topology Generator | - | str (path) | Path to AMBER coordinate (.rst7) file |
| selected_standard_forcefields | TLeap Topology Generator, Structure | - | dict | Selected standard force fields for tLEaP |

| | | | | |
|---|---|---|---|---|
| | Preprocessor (pipeline) | | | |
| single_state_ff_requirements | TLeap Topology Generator | - | dict | Force field requirements for single-state topology |
| single_state_selected_forcefields | TLeap Topology Generator | - | dict | Selected force fields for single-state topology |
| small_molecules | Small Molecule Parameterizer | - | list | Detected small molecules for parameterization |
| solvation_parameters | TLeap Topology Generator, Structure Preprocessor (pipeline) | - | dict | Solvation model parameters (explicit/implicit) |
| specific_capacitance | Protonation State Analyzer | - | float | Specific capacitance from protonation analysis |
| structure | PDB Loader | AltLoc Selector, Disulfide Bond Detector, MD Restraint Manager, Small Molecule Parameterizer | Structure | Current BioPython Structure (deprecated, backward compat) |
| structure_pdb_file | Structure Preprocessor (pipeline) | - | str (path) | Input PDB file for preprocessing pipeline |
| textbook_pkas | Protonation State Analyzer | - | dict | Textbook pKa values used as reference |
| tleap_input_file | TLeap Topology Generator | - | str (path) | Path to generated tLEaP input file |
| tleap_parameters | TLeap Topology Generator | - | dict | tLEaP execution parameters |

| | | | | |
|---|---|---|---|---|
| tleap_template | TLeap Topology Generator | - | dict | tLEaP input template for single state |
| topology | - | Molecular Dynamics Manager | str (path) | Path to topology file (required by MD Manager) |
| topology_extracted_pdb | Molecular Dynamics Manager | - | str (path) | PDB extracted from topology for visualization |
| transformed_pdb_file | Redox Site Preparer | - | str (path) | Path to redox-transformed PDB file |
| transformed_structure | Redox Site Preparer | - | Structure | Structure after redox transformation |
| transformer_info | Redox Site Preparer | - | dict | Redox transformer metadata |
| untransformed_structure | Redox Site Preparer | - | Structure | Structure before redox transformation |
| user_residue_classifications | Forcefield Parameterizer | - | dict | User classifications for non-standard residues |

**Supporting Information: PDB Record Types Parsed by ProPrep**

**Table S2. Header Record Types.** ProPrep parses 29 PDB header record types, presented to the user in categorized menus for targeted inspection. The table below lists each record type, the PDB format specification it follows, and the menu category under which it is displayed.

| Record Type | Description | Menu Category |
|---|---|---|
| HEADER | Classification, deposition date, PDB ID | Summary; Header Records |
| TITLE | Structure title | Summary; Header Records |
| AUTHOR | Authors | Header Records |
| COMPND | Compound (molecular) information | Header Records |
| SOURCE | Biological source organism | Summary; Header Records |
| KEYWDS | Keywords | Header Records |
| EXPDTA | Experimental technique | Summary; Header Records |
| REVDAT | Revision history | Header Records |
| CAVEAT | Severe warnings about the entry | Safety & Status Records |
| OBSLTE | Obsolete entry notification | Safety & Status Records |
| SPRSDE | Supersedes previous entry | Safety & Status Records |
| SPLIT | Split entry information | Safety & Status Records |
| HELIX | Helix secondary structure | Secondary Structure |
| SHEET | Beta-sheet secondary structure | Secondary Structure |
| SSBOND | Disulfide bond connectivity | Secondary Structure |
| LINK | Inter-residue linkages | Secondary Structure |
| SEQRES | Sequence residues per chain | Sequence Information |
| MODRES | Modified residue descriptions | Sequence Information |
| DBREF | Database cross-references (UniProt, GenBank) | Database References |
| SEQADV | Sequence differences and conflicts | Database References |
| CISPEP | Cis peptide bonds | Database References |
| JRNL | Journal citation information | Citations |
| HET | Heteroatom (non-standard residue) identifiers | Hetero Compounds |
| HETNAM | Heteroatom chemical names | Hetero Compounds |
| FORMUL | Molecular formulae for heterogens | Hetero Compounds |
| CRYST1 | Unit cell parameters and space group | Crystallographic Information |
| ORIGX | Coordinate transformation to original frame | Crystallographic Information |
| SCALE | Fractional-to-orthogonal transformation | Crystallographic Information |
| SITE | Active sites, binding sites, metal centers | Site Information |

**Table S3. REMARK Record Types.** ProPrep parses 41 REMARK record types, accessible through the Remark Records menu. Each REMARK number corresponds to a specific category of annotation defined by the PDB format specification.

| REMARK | Description |
|---|---|
| 0 | Re-refinement notice |
| 1 | Related publications |
| 2 | Resolution (Å) |
| 3 | Refinement statistics |
| 4 | Version and compliance |
| 100 | Processing site details |
| 200 | X-ray diffraction experiment |
| 205 | Fiber diffraction details |
| 210 | NMR experiment details |
| 215 | Solution NMR details |
| 217 | Solid-state NMR details |
| 230 | Neutron diffraction details |
| 240 | Electron crystallography |
| 245 | Electron microscopy (EM) |
| 247 | EM coordinate generation |
| 250 | Other experimental techniques |
| 265 | Solution scattering |
| 280 | Crystal properties (Matthews coefficient, solvent content) |
| 285 | CRYST1 unit cell details |
| 290 | Crystallographic symmetry operators |
| 300 | Biomolecule description |
| 350 | Biomolecule generation instructions |
| 375 | Special position atoms |
| 400 | Macromolecular compound details |
| 450 | Biological source details |
| 465 | Missing residues |
| 470 | Missing atoms |
| 475 | Zero-occupancy residues |
| 480 | Zero-occupancy atoms |
| 500 | Geometry and stereochemistry issues |
| 525 | Solvent atom distance information |
| 600 | Heterogen details |
| 610 | Missing atoms in heterogens |
| 615 | Zero-occupancy atoms in heterogens |
| 620 | Metal coordination |
| 630 | Inhibitor/compound description |
| 650 | Helix details |
| 700 | Sheet details |
| 800 | Important functional sites |
| 900 | Related PDB entries |

### Supporting Information: Water Analysis Methods

### Table S4: Water Analysis Methods in ProPrep's PDB Filter Module

ProPrep provides seven analysis methods for discriminating structurally important water molecules from bulk solvent. Each method can be applied individually or in combination, and parameters are user-configurable.

---

#### 1. Metal Ion Coordination

**Purpose:** Identify water molecules coordinating metal ions.

ProPrep identifies all metal atoms in the structure and computes the distance from each water oxygen to the nearest metal. Waters within the coordination distance are flagged as metal-coordinating.

| Parameter | Default | Description |
|---|---|---|
| Metal distance cutoff | 2.5 Å | Maximum distance for metal coordination |

**Reported metrics:** Nearest metal identity, metal–oxygen distance, coordination flag.

---

#### 2. Hydrogen-Bonding Network Analysis

**Purpose:** Quantify and characterize hydrogen-bonding interactions around each water molecule.

For each water, ProPrep searches for potential hydrogen-bond partners among protein, ligand, and other water atoms within the distance cutoff. Candidate bonds are evaluated using a combined distance–angle scoring function. The heavy-atom angle at the shared hydrogen position is estimated, and scores are assigned as follows: angles ≥150° receive a score of 1.0, ≥120° a score of 0.8, ≥90° a score of 0.6, and <90° a score of 0.3. The distance score is computed as 1.0 − (distance / cutoff). The combined score is the product of the distance and angle scores. Up to the maximum number of hydrogen bonds are accepted per water, ranked by combined score.

| Parameter | Default | Description |
|---|---|---|
| H-bond distance cutoff | 3.5 Å | Maximum donor–acceptor distance |
| H-bond atoms | N, O, S | Element types eligible for H-bonding |
| Max H-bonds per water | 4 | Maximum accepted H-bonds per water |

**Reported metrics:** H-bond count by partner type (protein, water, hetero), total count, partner identities with distances and angle scores.

---

#### 3. B-Factor Analysis

**Purpose:** Assess water mobility from crystallographic temperature factors.

The B-factor (temperature factor) for each water oxygen is extracted directly from the PDB file. Low B-factors indicate well-ordered, structurally significant waters; high B-factors suggest mobile waters that are less likely to be structurally important.

**Reported metrics:** B-factor value for the water oxygen atom.

---

#### 4. Burial Analysis

**Purpose:** Quantify how deeply a water molecule is embedded within the protein.

Burial is assessed in three phases of increasing detail.

*Phase 1: Proximity-based burial.* All atoms within a search radius around the water are counted, with counts separated by type (protein, hetero, water, metal). Three weighting schemes are available: (a) uniform counting (each atom = 1.0), (b) distance-weighted (weight = 1/distance), and (c) van der Waals-weighted (weight = sum of vdW radii / distance). The burial percentage is computed as the ratio of the total weight to a configurable reference value for full burial.

| Parameter | Default | Description |
|---|---|---|
| Burial radius | 5.0 Å | Search radius around water |
| Burial atom types | protein, hetero | Which residue types to count |
| Burial weighting | count | Weighting scheme (count, distance, vdw) |
| Burial max expected | 50 | Reference atom count for 100% burial |

*Phase 2: Multi-radius burial profiling.* Burial is computed at multiple radii from 2.0 to 8.0 Å in 0.5 Å steps, generating a burial-versus-radius profile. The profile reveals the saturation radius (where burial increases by <10% with further radius expansion) and the radius range of steepest burial increase. Results are displayed as ASCII charts in the terminal.

| Parameter | Default | Description |
|---|---|---|
| Minimum radius | 2.0 Å | Smallest search radius |
| Maximum radius | 8.0 Å | Largest search radius |
| Step size | 0.5 Å | Radius increment |

*Phase 3: Directional burial analysis.* Three-dimensional space around the water is divided into 8 compass sectors (N, NE, E, SE, S, SW, W, NW) using the azimuthal angle. Atoms within the burial radius are assigned to sectors and weighted. The analysis identifies the primary burial direction (most shielded), secondary direction, and pocket opening (least shielded direction, representing the water's potential exit route). Burial patterns are classified as uniform, moderately directional, or highly directional based on the range of sector weights relative to the mean.

**Reported metrics (all phases):** Burial percentage, raw atom count, atom counts by type, closest atom distance, estimated SASA, saturation radius, burial profile, sector weights, primary/secondary burial directions, pocket opening direction, pattern classification.

---

#### 5. Protein-Protein Interface Proximity

**Purpose:** Identify waters bridging multi-chain interfaces.

For structures with multiple protein chains, ProPrep determines whether atoms from two or more different chains are within the interface distance cutoff of each water. Such waters may mediate inter-chain contacts and should generally be retained.

| Parameter | Default | Description |
|---|---|---|
| Interface distance cutoff | 5.0 Å | Maximum distance to count as interface proximity |

**Reported metrics:** Interface flag (boolean), identities of bridged chains.

---

#### 6. Solvent-Accessible Surface Area (SASA)

**Purpose:** Quantify the solvent exposure of each water molecule.

SASA is computed for the entire structure using the Lee-Richards algorithm via the FreeSASA library (if installed), with a probe radius of 1.4 Å. Low SASA values indicate buried waters. If FreeSASA is unavailable, SASA is estimated from the burial analysis (estimated SASA = 40.0 × (1 − burial_percentage / 100)).

| Parameter | Default | Description |
|---|---|---|
| Probe radius | 1.4 Å | Solvent probe size for SASA calculation |

**Reported metrics:** SASA value ($Å^2$).

---

#### 7. Water Network Analysis

**Purpose:** Characterize the topology of hydrogen-bonded water networks.

ProPrep constructs a graph (using the NetworkX library) where water molecules are nodes and hydrogen bonds between waters are edges. The analysis identifies: (a) connected components (clusters of hydrogen-bonded waters), (b) hub waters with three or more hydrogen-bonded water neighbors, (c) the longest water chains (diameter paths through tree-like clusters), and (d) isolated waters with no water–water hydrogen bonds.

| Parameter | Default | Description |
|---|---|---|
| Network type | water_only | Restrict network to water–water H-bonds |
| Minimum cluster size | 2 | Smallest cluster reported |

| | | |
|---|---|---|
| Show isolated | true | Whether to list isolated waters |

**Reported metrics:** Total connections, number of connected components, significant cluster count and sizes, hub water identities and degrees, longest chain lengths and paths, isolated water identities.

---

**Table S5: Water Categorization**

Based on the analysis results, each water molecule is assigned to one of the following categories (in priority order):

| Category | Criteria |
|---|---|
| Metal-coordinating | Within metal distance cutoff of a metal ion |
| Highly connected | ≥3 total hydrogen bonds |
| Interface | Bridges atoms from 2+ protein chains |
| Buried | Burial percentage above threshold |
| Ordered | B-factor < 30 Å² |
| Bulk solvent | Does not meet any of the above criteria |

Results for all waters are displayed in a color-coded table sorted by category priority. Users interactively select which waters to retain based on the analysis results and, optionally, visual inspection in the 3D structure viewer.

---
